# Dynamic Ribosomal RNA Methylation Regulates Translation in the Hematopoietic System and is Essential for Stem Cell Fitness

**DOI:** 10.1101/2025.09.29.679281

**Authors:** Ofri Rabany, Sivan Ben Dror, Maram Arafat, Hadar Aharoni, Yudit Halperin, Virginie Marchand, Nikolai Romanovski, Noga Ussishkin, Maayan Livneh, Adi Reches, Judith Wexler, Nina Mayorek, Galia Monderer-Rothkoff, Sagiv Shifman, Widad Mammer Bouhou, Michael VanInsberghe, Cornelius Pauli, Carsten Müller-Tidow, Ola Karmi, Yoav Livneh, Alexander van Oudenaarden, Yuri Motorin, Daphna Nachmani

## Abstract

Self-renewal and differentiation are at the basis of hematopoiesis. While it is known that tight regulation of translation is vital for hematopoietic stem cells’ (HSCs) biology, the mechanisms underlying translation regulation across the hematopoietic system remain obscure. Here we reveal a novel mechanism of translation regulation in the hematopoietic hierarchy, which is mediated by ribosomal RNA (rRNA) methylation dynamics.

Using ultra-low input ribosome-profiling, we characterized cell-type-specific translation capacity during erythroid differentiation. We found that translation efficiency changes progressively with differentiation and can distinguish between discrete cell populations as well as to define differentiation trajectories. To reveal the underlying mechanism, we performed comprehensive mapping of the most abundant rRNA modification - 2’-O-methyl (2’OMe). We found that, like translation efficiency, 2’OMe dynamics followed a distinct trajectory during erythroid differentiation.

Genetic perturbation of individual 2’OMe sites demonstrated their distinct roles in modulating proliferation and differentiation. By combining CRISPR screening, molecular and functional analyses, we identified a specific methylation site, 28S-Gm4588, which is progressively lost during differentiation, as a key regulator of HSC self-renewal. We showed that low methylation at this site led to translational skewing, mediated mainly by codon frequency, which promoted differentiation. Functionally, HSCs with diminished 28S-Gm4588 methylation exhibited impaired self-renewal capacity *ex-vivo*, and loss of fitness *in-vivo* in bone marrow transplantations.

Extending our findings beyond the hematopoietic system, we also found distinct dynamics of 2’OMe profiles during differentiation of non-hematopoietic stem cells. Our findings reveal rRNA methylation dynamics as a general mechanism for cell-type-specific translation, required for cell function and differentiation.

**KEY POINTS:**

1. Hematopoietic differentiation is associated with rRNA methylation dynamics to control cell-type-specific translation.
2. Translation efficiency can distinguish discrete cell types and define differentiation trajectories.
3. HSC fitness is regulated by a single rRNA methylation.

## INTRODUCTION

Multi-cellular organisms rely on cellular differentiation and acquisition of cell identity for their proper development. Hematopoietic stem cells (HSCs) are at the apex of the hematopoietic hierarchy, and sustain lifelong blood production by balancing self-renewal with differentiation^1^. Previous studies demonstrated that HSCs have a tightly regulated translation rate, and when aberrant leads HSCs to exit their quiescent state resulting in impaired function^2–10^. The importance of translation regulation in HSCs can be further appreciated from studies of a group of inherited human syndromes known as ribosomopathies, which are caused by mutations in ribosomal or ribosome-associated proteins. Ribosomopathies commonly present with bone marrow failure and a strong predisposition for leukemia development^11–16^. Interestingly, these syndromes often exhibit cell-type-specific hematopoietic features - such as erythroid and megakaryocytic defects in Diamond-Blackfan anemia^13,18,19^, while neutropenia is common in Shwachman-Diamond syndrome^12,17^. This selectiveness of pathological features might suggest a role for cell-type-specific translation in the hematopoietic system.

In addition to ribosomal proteins, each ribosome also carries more than 250 chemically modified rRNA nucleotides. The most prevalent rRNA modification is 2’-O-methylation (2’OMe) of the ribose, required for proper processing and folding of rRNAs, and hence is crucial for translation^18–23^. Each 2’OMe site is installed by a specific C/D box small nucleolar RNA (snoRNA/snoRD), which guides the methylation machinery to the correct rRNA site via extensive base-pairing. This long base-pairing results in an almost exclusive relationship between a snoRNA and a modified site, where usually a snoRNA will facilitate the modification of a single site^18,24^. Previous work by us and others has shown that global perturbation to rRNA modifications, and specifically to 2’OMe, leads to aberrant hematopoiesis and to leukemia susceptibility^25–28^, which are key features of ribosomopathies. As 2’OMe is a modulator of translation^20,22^, known to be at sub-stoichiometric levels^29–31^, we hypothesized that changes to 2’OMe levels across the hematopoietic hierarchy, can lead to cell-type-specific translational capacities with important implications for physiology.

To address this, we compared the transcriptome, translatome, and rRNA methylome of rare populations along erythroid differentiation, enabled by recent advances in ultra-low input ribosome profiling^32^ and comprehensive mapping of the 2’OMe landscape^33^. Our analyses revealed dynamics of translation capacities and of 2’OMe levels during hematopoietic differentiation. Importantly, extending beyond hematopoiesis, we also observed distinct patterns of 2’OMe dynamics during non-hematopoietic differentiation, suggesting that rRNA modification profiles are tailored to support cell identity across different biological systems.

## METHODS

### Ultra-low input ribosome profiling

Ultra-low ribosome profiling was performed by FACS sorting 1000 cells directly into cooled lysis buffer. Ribosome profiling libraries were prepared using a version of the scRibo-seq protocol^32^ that was adapted for higher cell inputs (1000 cells), to achieve this, we scaled up all reaction volumes 100×. Ribo-seq raw reads were processed, aligned, and quantified as described in scRibo-seq^32^. Please see supplemental method for further details.

### Mouse HSPCs isolation and transplantation

Bones from 8-week-old mice were collected. Cells were ACK treated and washed with PBS+2%FBS. Lineage^+^ cells were depleted by mouse lineage-depletion kit (Miltenyi Biotec), and sorted on a AriaIII BD sorter. In BMT C57BL/6J recipient mice were pre-treated with Baytril then irradiated by two doses of 550 cGy. Please see supplemental method for further details.

## RESULTS

### Ultra-low ribosome profiling reveals progressive translational changes during erythroid differentiation

To explore translation modulation in the hematopoietic system, we focused on mouse erythroid differentiation as this lineage is commonly affected in ribosomopathies^11,12^. We performed ultra-low ribosome profiling, which included both ultra-low RNA-seq and ultra-low sequencing of ribosome-protected fragments (RFP), of 5 populations – hematopoietic stem cells (HSC), preMegE (megakaryocytes and erythrocyte progenitors), MkP (megakaryocytes progenitors), preCFUE and CFUE (both erythrocyte specific) populations (**Fig.1A**). We FACS sorted 10,000 cells of the above-mentioned populations, for RNA-seq (**Fig.S1A** and **S1B, Supplementary Table 1**), and from the same samples we further sorted 1,000 cells for the processing and sequencing of RPFs (**Fig.1B**, and **supplementary Table 2,** QC in **Fig.S1C**). This unique experimental setup allows us to assess the different qualities of the RNA-seq compared to the RPFs, as well as to calculate translation efficiency (TE), a measure of RPF normalized to transcript levels^34^. Thus, our ultra-low ribosome profiling enables us to evaluate unique translational capacities of each population.

**Figure 1.**
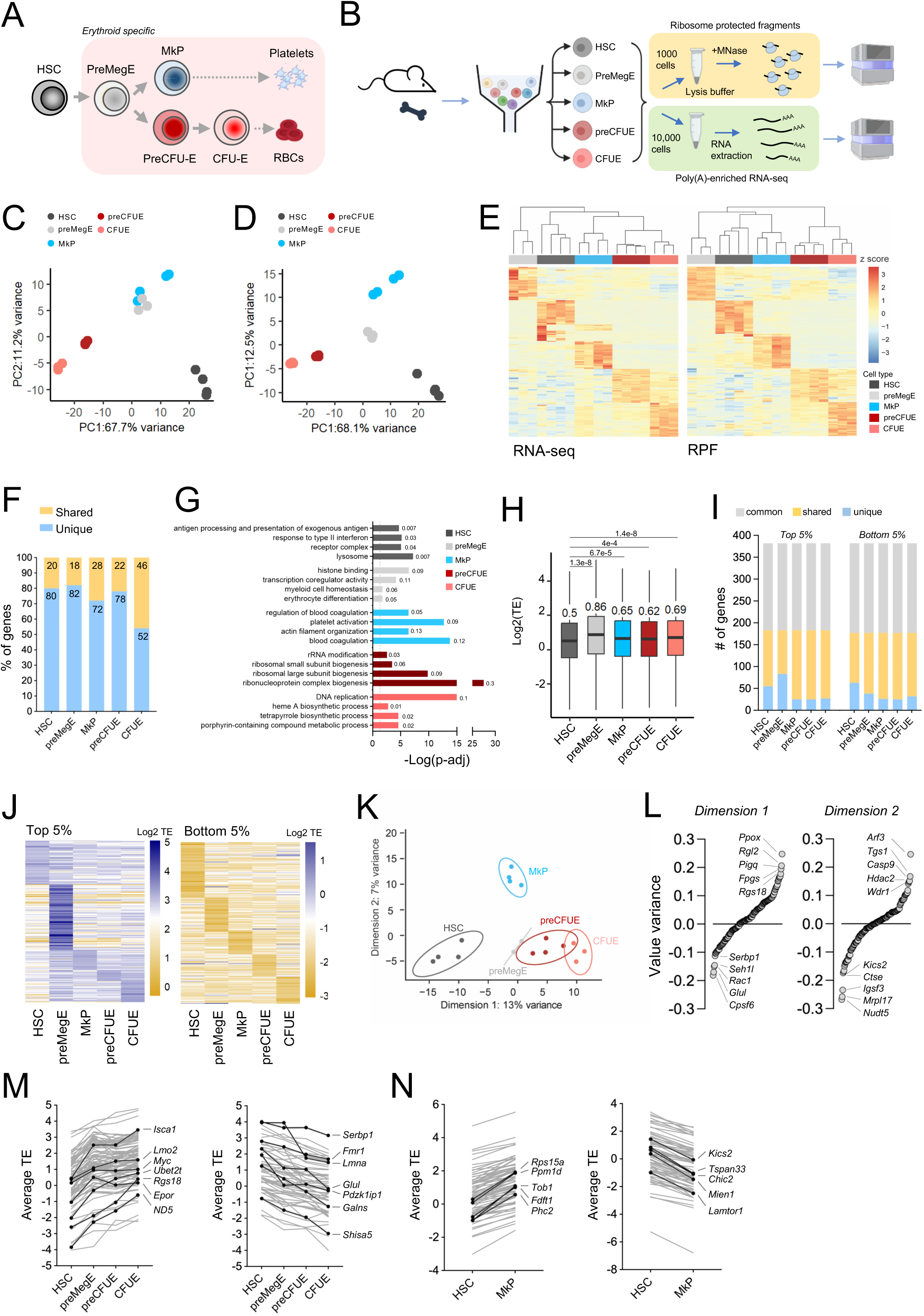
Ultra-low ribosome profiling of erythroid differentiation. (A) Schematic illustration of the relevant populations in the hematopoietic hierarchy. Hematopoietic stem cells (HSC) are at the apex, preMegE are erythroid committed progenitors, megakaryocyte progenitors (MkP) are committed to generate megakaryocytes (platelets), preCFUE and CFUE are erythrocyte-committed. Platelets and red blood cells (RBCs) are depicted as the terminally differentiated cells but were not analyzed. HSCs were sorted using the following markers Lin^-^Sca1^+^cKit^+^CD48^-^CD150^+^, preMegE – Lin^-^cKit^+^Sca1^-^Il7R^-^CD41^-^CD16/32^-^CD150^+^CD105^-^, preCFUE - Lin^-^cKit^+^Sca1^-^Il7R^-^CD41^-^CD16/32^-^CD150^+^CD105^+^, CFUE - Lin^-^cKit^+^Sca1^-^Il7R^-^CD41^-^ CD16/32^-^CD150^-^CD105^+^, MkP - Lin^-^cKit^+^Sca1^-^Il7R^-^CD41^+^ .(B) Schematic illustration of experimental setup of ultra-low ribo-seq (sequencing of ribosome protected fragments, top) and RNA-seq (bottom). For ultra-low ribo-seq analysis 1,000 cells were sorted and for ultra-low RNA-seq 10,000 cells from the same sample were sorted. 3-4 biological replicates were made for each cell type, each replicate consisted of pooled 8 mice. (C and D) PCA analysis of the top 500 variance genes of DESeq variance stabilizing transformed (VST) counts of ultra-low RNA-seq (C) and of RPF generated by ultra-low ribo-seq (D). (E) Heat map generated from unsupervised clustering of 50 signature genes for ultra-low RNA-seq (left) and for ultra-low ribo-seq (right). Signature genes were defined as the topmost highly expressed genes (according to log2FC) in a specific cell compared to all other cell types, and with FDR<0.05. (F) The percentage of shared (gold) and unique (blue) genes between the signature genes identified for ultra-low RNA-seq and ultra-low ribo-seq in E. (G) GO analysis on all significant signature genes (identified as in E) of RPF dataset (ribo-seq). The small numbers next to the bars are gene ratios. Number of genes in each group: HCS = 1476, PreMegE = 234, MkP = 201, PreCFUE = 147, CFUE = 526. (H) Global TE in specific cell types. Statistical significance was calculated by Welch ANOVA (p=1.76e-31) and Games-Howell post-hoc test (plot shows p. Adjusted values comparing to HSCs). (I) Focusing on the top or bottom 5% of TE genes, we categorize 3 categories – genes which are shared by all cell types (gray), that are shared by several but not all cell types (gold), and those which are uniquely in the 5% of a single cell type (blue). (J) Heat map of the unique TE genes of the top 5% (high TE, left) or bottom 5% (low TE, right) according to cell type. (K) Sparse PLS discriminant analysis (sPLS-DA), a supervised clustering model, of TE in the various populations. By applying sPLS-DA on TE values we identify 150 genes that enable discrimination of cells on the x-axis (Dimension 1) and 150 genes on the y-axis (Dimension 2). (L) Ranking according to value variance of dimension 1 (left) and dimension 2 (right) of sPLS-DA in K. Specific top 5 genes are depicted. (M) Shown are the progressive changes to TE in genes identified in sPLS-DA of dimension 1 (x-axis). Divided into progressively increasing TE genes (left) and progressively decreasing TE genes (right). In black are specific examples of genes with possible relevant roles. (N) Shown are the progressive changes to TE in genes identified in sPLS-DA of dimension 2 (y-axis). Divided into progressively increasing TE genes (left) and progressively decreasing TE genes (right). In black are specific examples.

We initially analyzed each dataset separately, principal-component analysis (PCA) using variance stabilizing transformed counts (VST) of the ultra-low RNA-seq data showed some clustering of the populations, however, preMegE and MkP clustered together even though they are distinct populations^35^ (**Fig.1C**). In contrast, PCA of RPF’s VST counts showed distinct clustering of each population including discrimination between preMegE and MkP (**Fig.1D**). We found that RPF data clusters the populations better than RNA-seq data also when using PC2, PC3 and PCA analysis with TPM values, showing the robustness of RPF ability to cluster the populations (**Fig.S1D**). Of the variable genes used for PCA, almost half of the genes (212 genes out of 500, 46%), were non-overlapping and exclusive to each data set, suggesting that RFP data offers new insight into these populations. Unsupervised hierarchical clustering of the cells’ signature genes generated unique fingerprints for each cell type both in the RNA-seq and the RPF datasets (**Fig.1E**). Notably, the signature genes of the two datasets shared only 15-46% overlap, depending on the specific population (**Fig.1F**). Among the signature genes detected exclusively in the RPF datasets are *Mecom* and *Flt3* in HSC, both with well-established roles in HSCs biology and leukemia development^36–39^ (**Supplementary Table 3**). Additional examples include *Vwf* and *Itga2,* known markers of MkP^40–42^, and *Hebp1* in CFU-E^43,44^ (**Supplementary Table 3**). Gene Ontology (GO) analysis of RPF signature genes revealed cell type-specific pathway enrichments associated with relevant biological functions (**Fig.1G**).

Next, we combined the two datasets to calculate TE for each population. We found that HSCs have the lowest global TE levels, while preMegE have the highest TE of the tested populations (**Fig.1H** and **Fig.S1E**). Thus confirming by ribosome profiling previous reports of low translation rate in HSCs and high translation rate in progenitors^8^. By examining the top and bottom gene at the 5% of TE (**Fig.S1F**), we identified genes that are common across all populations, shared among subsets, and, importantly, distinct to each cell type (**Fig.1I-J**). Interestingly, in the high TE genes found exclusively in HSCs we found chromatin modulators such as *Dnmt3a* and *Kdm3b,* with known roles in HSC biology^45–51^. In this group we also identified *Tsc1*, a mTOR inhibitor crucial for HSC biology^52,53^, and *Nop58* which is involved in ribosome biogenesis and rRNA methylation^54,55^ (**Supplementary Table 4**). In the low TE genes found exclusively in HSCs we find *Eif2s2*, a translation initiation factor, suggesting it might play a role in translation regulation. We were able to identify unique genes for each population even when using a permissive threshold of top or bottom 10% and 20% of TE (**Fig. S1G**). This emphasized that TE, as a measure of translation, carries cell-type-specific characteristics.

By applying a supervised clustering model, we found that TE can be used to separate cell types according to their differentiation trajectory (**Fig.1K**). This analysis identified 150 genes, in which TE levels can discriminate populations on each clustering axis. Interestingly, the first clustering dimension mainly separated the populations according to erythrocyte development (HSC-preMegE-preCFUE-CFUE), whereas the second dimension separated the megakaryocyte progenitors, MkP, from all other populations (**Fig.1K-L**). In dimension 1 we found genes with progressively increased or decreased TE (**Fig.1M**). In the increasing TE genes group we found *Epor* and *Myc* (essential for erythropoiesis ^56,57^), and in the decreasing group we found *Fmr1* and *Lmna,* with known roles in human diseases^58–60^ and suggested roles in regulating hematopoietic cell biology^61–63^. In dimension 2, we found additional genes involved in hematopoietic physiology - *Ppm1d*, a regulator of p53 known to have a role in clonal hematopoiesis^64^, *Chic2,* which is deleted or translocated in leukemia^65–67^, and the mTOR regulators – *Lamtor1* and *Kics2* (**Fig.1N**).

In summary, while it is well-established that HSCs have a unique translational capacity, here we used ultra-low ribosome profiling to show that this is not exclusive to HSCs. Instead, our data reveal an intricate picture, in which TE ranking can provide a discrete, cell-type-specific signature while TE dynamics can be a continuous trait, exhibiting progressive changes in translational capacity along differentiation trajectories.

### Mapping of rRNA 2’OMe profiles in erythroid differentiation reveals progressive methylation dynamics

Translational control is essential for HSC function and hematopoiesis^8^. Although rRNA modifications are known to influence translational capacity, their specific contributions to the regulation of physiological processes remain largely unexplored. As rRNA methylation can occur at sub-stoichiometric levels^29–31^, we hypothesized that changes to rRNA 2’OMe profiles might have a role in translation regulation impacting cellular function. To this end, we comprehensively mapped 2’OMe landscape during erythroid differentiation (**Fig.1A**). We again sorted HSCs, preMegE, MkP, preCFUE and CFUE populations, extracted total RNA and performed RiboMethSeq^33^ to quantify 2’OMe levels on rRNAs, at single nucleotide resolution (**Fig.2A**). These analyses revealed variable positions in which not all nucleotides are modified (*i.e.* MethScore is less than 1, **Fig.2A**). This is in accordance with previous findings of changes to rRNA modifications in other biological systems^29,68–71^, suggesting that rRNA methylation levels are also actively modulated in the hematopoietic system. Specifically, we found 2’OMe dynamics in the erythrocyte differentiation path from HSC and preMegE, through preCFU-E to CFU-E, but less significant dynamics along the megakaryocytic path from HSC to MkP (**Fig.2A-C, Supplementary Table 5**). This is evident by PCA and Euclidian distance analysis, where no differences were observed between preMegE and MkP (**Fig.2B-C**). Our findings thus suggest that the 2’OMe landscape is tightly regulated and that 2’OMe plasticity might have a role in regulating hematopoietic cells’ function and identity.

**Figure 2.**
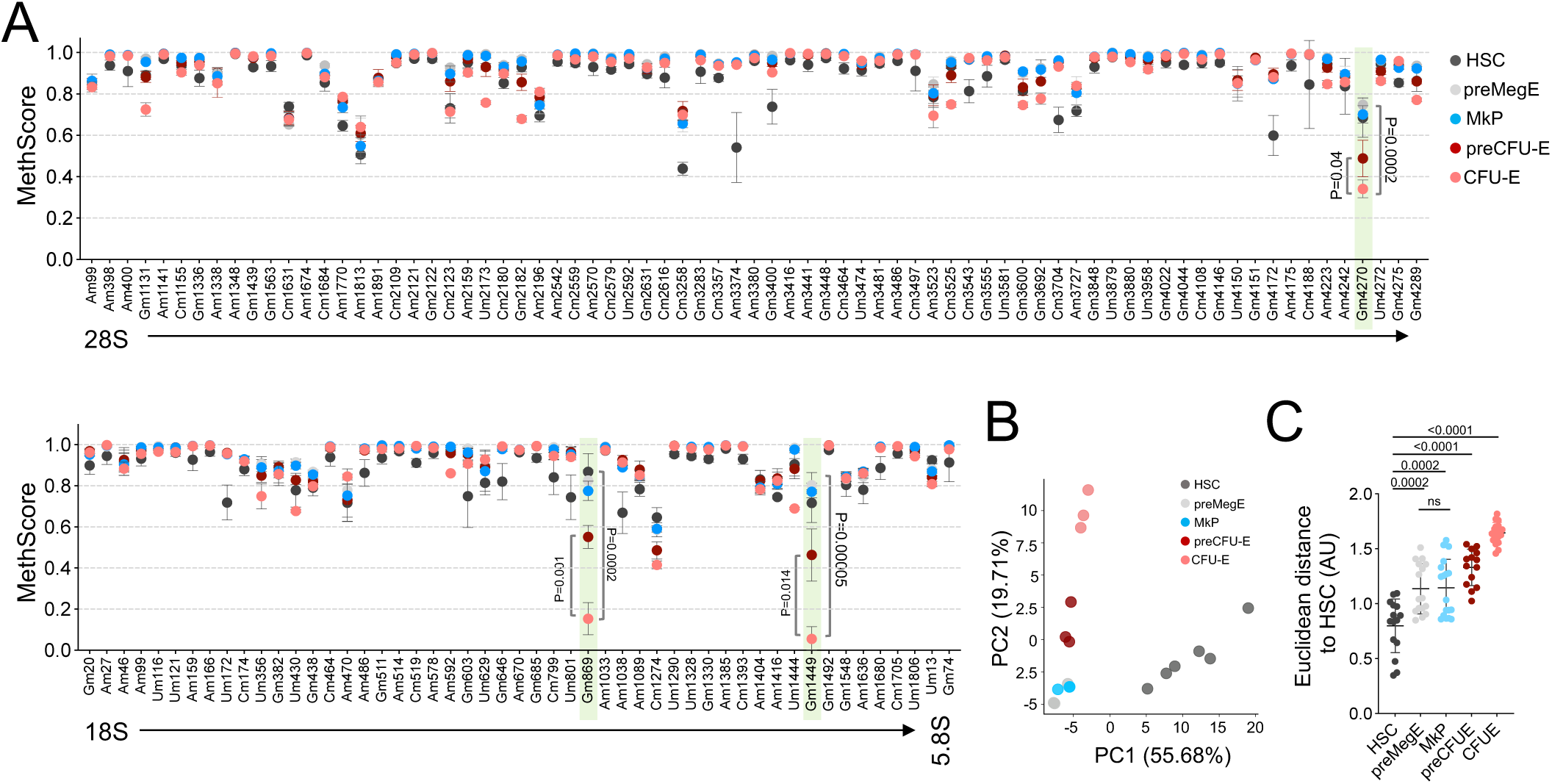
2’OMe dynamics in erythroid differentiation. (A) RiboMethSeq analysis of HSCs, preMegE, MkP, preCFUE and CFUE. Highlighted in green are the most variable modifications. MethScore is shown for known 28S, 18S and 5.8S sites (x-axis). A value of 1 represents a fully modified nucleotide, 0 is unmodified. Data are mean±SD of 4 biological replicates, each replicate consists of a pooling of 4 mice. Significance was determined by t-test. P-values of all known methylation sites are listed in Supplementary Table 5. (B) PCA of HSC, preMegE, MkP, preCFU and CFUE using methscore. (C) Euclidean distance analysis to quantitively evaluated dissimilarities of 2’OMe profiles of the indicated cell-type calculated relative to HSC. The Euclidean distance represents dissimilarity, hence the higher value, the less similar is the specific cell-type to HSC. Significance was determined by t-test.

### Functional analyses demonstrate single 2’OMe sites with differential regulatory capacity of proliferation and erythrocyte differentiation

Next, we aimed to demonstrate the functionality of 2’OMe dynamics on cellular traits and translation. We chose to focus our analyses on the 3 main sites that showed the highest dynamics – 18S-Gm869, 18S-Gm1449 and 28S-Gm4270 (**Fig.2A**, highlighted bars and **Fig.S2A**). 2ʹOMe sites are highly conserved across evolution, so the mouse 2’OMe sites have corresponding human sites that are also modified^72,73^. In this manuscript, while we will alternate between mouse and human cells, we will use from now on the human nomenclature for convenience and consistency with previous publications (**Table 1**).

**Table 1.**
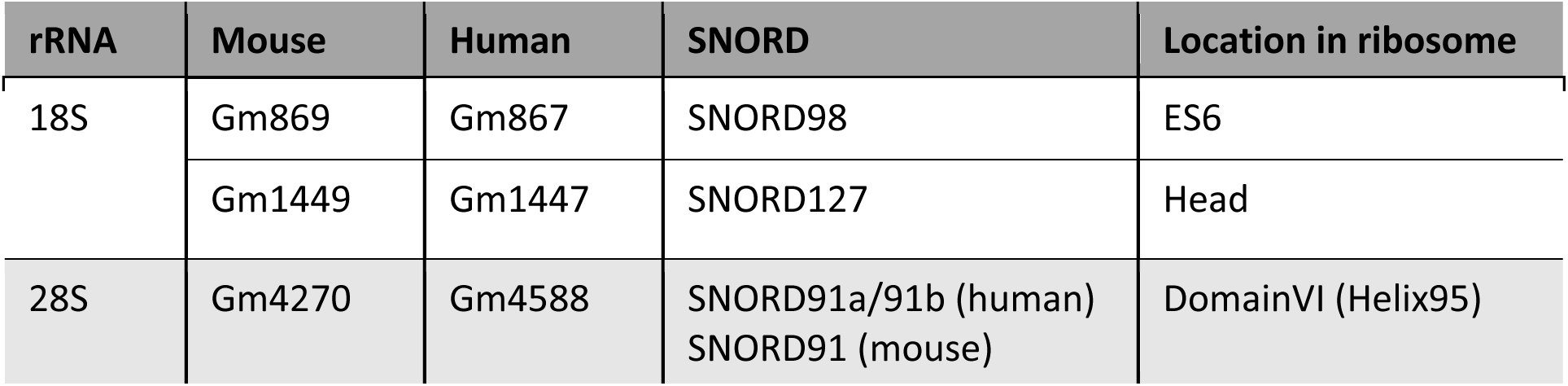
Nomenclature of the studied mouse and human rRNA methylation.

We mapped the three variable 2ʹOMes onto the ribosome structure and found them positioned at functionally important sites^74^ with potential roles in regulating translation and cell properties (**Fig.3A**). Specifically, human 18S-Gm867 (mouse 18S-Gm869) resides in expansion segment 6 (ES6), previously implicated in the binding of initiation factors eIF3e and eIF4A^75,76^. Human 18S-Gm1447 (mouse 18S-Gm1449) is located in the “head” of the small subunit, a region involved in ribosome translocation^77^. Finally, human 28S-Gm4588 (mouse 28S-Gm4270) lies within Helix 95, part of the highly conserved Sarcin/Ricin Loop, which is an integral component of the ribosome’s GTPase-associated center, important for binding initiation and elongation factors^78–80^.

**Figure 3.**
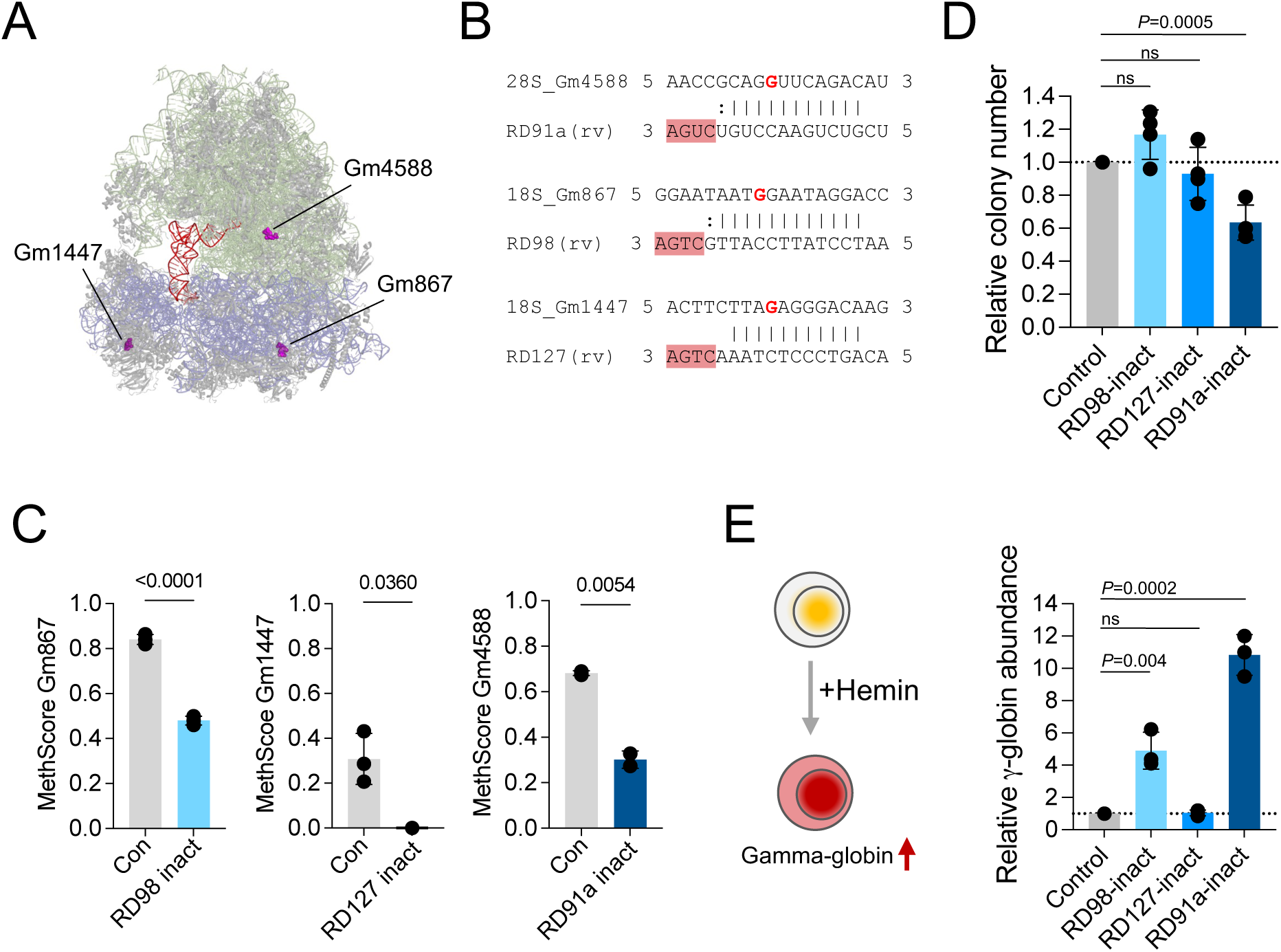
Specific 2’OMe sites show distinct capacities to regulate cellular proliferation and differentiation. (A) Location of the variable 2’OMe within the human ribosome structure (PDB 8glp). Light green – 28S rRNA, lilac – 18S rRNA, red – P-site, orange – mRNA, magenta – specific modified nucleotides, gray – ribosomal proteins. (B) Alignment between specific *SNORDs* and their corresponding methylation sites. Highlighted in dark pink are the snoRNA’s D’ box. In red is the modified nucleotide on rRNA. The snoRD sequence is presented in reverse nucleotide order (rv). (C) RiboMethSeq analysis of SNORD-specific inactivated K562 cells. Data are mean±SD of two or three biological replicates and show the MethScore of the specific modified site. (D) Colony formation unit assay to assess clonogenic growth of specific snoRD-inactivated K562 cells. Cells were plated in semi-solid media and number of colonies was assessed 7 days after plating. Data are colony number relative to the number of colonies in control in each biological repeat (represented by each black circle). Data are presented as mean±SD of 3-4 biological replicates, each performed in triplicates. Significance was determined by t-test. ns – not significant. Dotted line indicates 1 threshold for convenience. (E) qPCR analysis of gamma-globin levels after hemin-induced differentiation of SNORD-inactivated K562 cells relative to control cells. Data are mean±SD of three biological replicates, each performed in triplicates. Dotted line indicates 1 threshold for convenience. Statistically significant differences are indicated and determined by t-test. ns – not significant.

To causally link the three dynamic 2ʹOMe sites to cellular function, we identified their guiding snoRDs: *SNORD98* for 18S-Gm867, *SNORD127* for 18S-Gm1447, and *SNORD91a* and *SNORD91b* for 28S-Gm4588 (Fig.3B, Fig.S2B). Using CRISPR–Cas9 in K562 erythroid leukemia cells, we introduced inactivating mutations in the relevant snoRDs, resulting in reduced snoRD expression and corresponding decreases in 2ʹOMe levels (**Fig.S2C**, **Fig.3C**). Unfortunately we were unsuccessful in inactivating *SNORD91b*, leading us to focus only on SNORD91a-inact cells, which showed the expected ∼50% reduction of Gm4588 (**Fig.3C**). Since these snoRDs are processed from introns, we verified that the host genes remained intact and that predicted off-targets were unaffected (**Fig.S2D–E**). Interestingly, *Snord98* and *Snord127* levels in specific cell types do not follow the progressive dynamics of their respective methylation sites (**Fig.S2F**), supporting our hypothesis that 2’OMe levels are actively regulated.

Our functional analyses focused on two hallmark traits of hematopoietic cells - clonogenic growth and differentiation capacity. We observed distinct effects of snoRD inactivation on clonogenic growth. Inactivation of *SNORD127* or *SNORD98* had no impact, whereas *SNORD91a* inactivation significantly reduced colony numbers (**Fig.3D**), indicating that even a 50% reduction in Gm4588 has a functional implication. Upon hemin-induced erythroid differentiation^81^, *SNORD127* inactivation again showed no effect, while *SNORD98* or *SNORD91a* inactivation enhanced γ-globin levels (**Fig.3E**), reflecting increased responsiveness to differentiation agents. These findings demonstrate that individual methylation sites differentially regulate cellular properties. Importantly, they suggest that reduced Gm867 or Gm4588 methylation during differentiation (**Fig.2A**) may directly affect cellular function.

### CRISPR *SNORD*-screen identifies a role for 28S-Gm4588 during megakaryocytic differentiation

To gain additional insights into the possible role of 2’OMe in megakaryocyte development, we performed a CRISPR screen of snoRDs in a model of megakaryocyte differentiation (**Fig.4A**). We designed a CRISPR library targeting human snoRDs^26^, carrying a GFP cassette. We used K562 cells, which upon PMA treatment acquire megakaryocyte-like features^82^ - increased cell size and the expression of the megakaryocytic marker CD41/61, which is not expressed by K562 in steady state (**Fig.S3A**). In this screen, K562 cells were transduced with the CRISPR library, GFP^+^ cells were sorted, cultured, and were either left untreated or treated with PMA. Then CD41/61^high^ cells were sorted and analyzed for sgRNA enrichment (**Fig. 4A**, and **Fig.S3B**). To our surprise, our analysis found only a single snoRD which was significantly enriched in the differentiated cells – *SNORD91a* (**Fig.4B**). All other significant sgRNAs were depleted from the differentiated population, and mainly targeted control protein-coding genes; none were snoRD-specific sgRNAs (**Fig.4B** and **Fig.S3C**).

**Figure 4.**
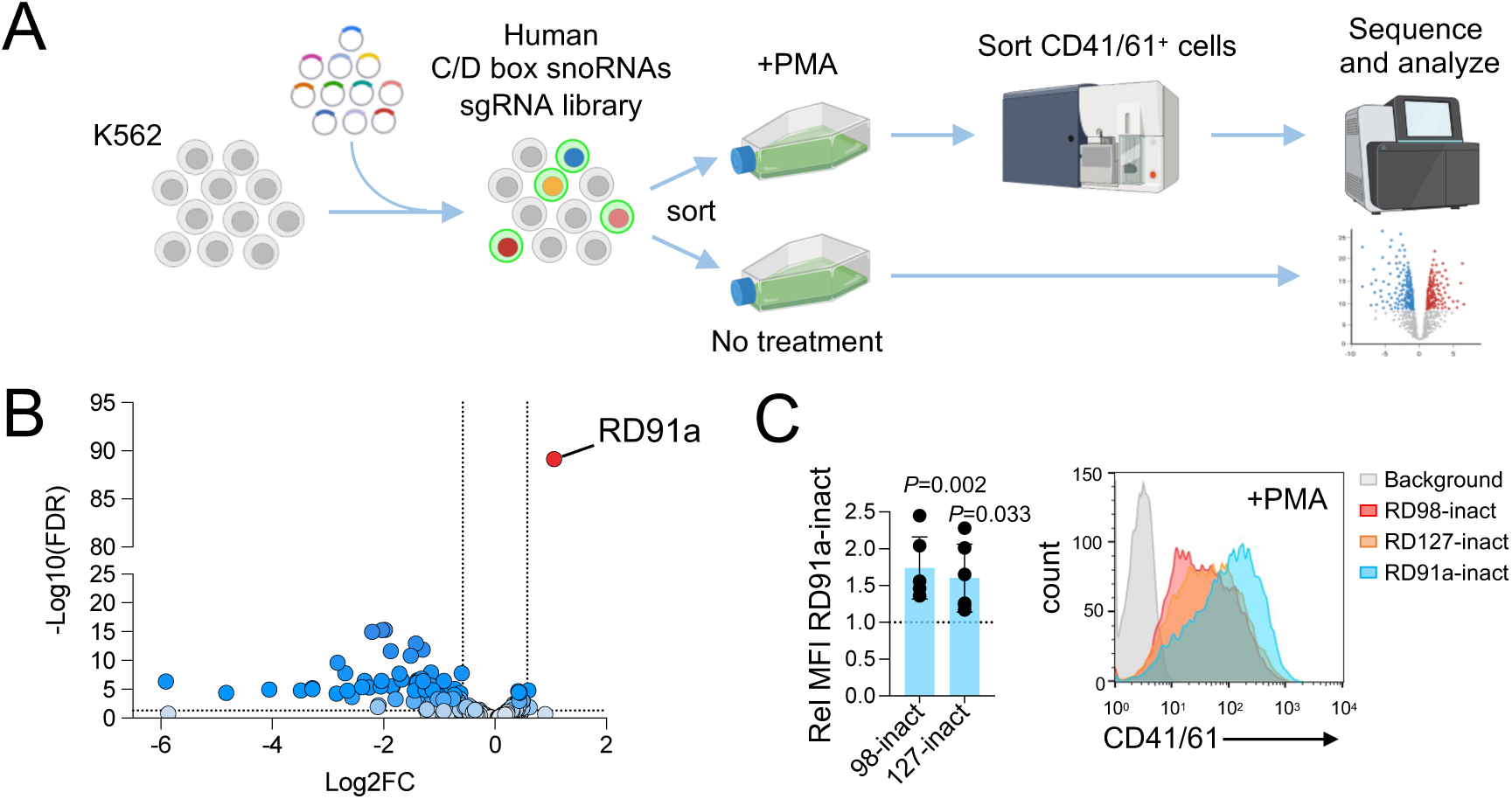
Human C/D box snoRNA CRISPR screen reveals a role for Gm4588 in megakaryocytic differentiation. (A) A scheme of CRISPR screen. K562 cells were transduced with human *SNORD*-targeting CRISPR library at a low MOI (=0.3). Two days after transduction GFP^+^ cells were sorted and cultured for 1 week prior to PMA treatment (2nM for 72hrs). Differentiated cells (CD41/61^high^ cells) were sorted from the PMA treated experimental group. The screen was performed in 4 independent replicates. (B) Volcano plot of sgRNA enrichment of SNORD CRISPR screen, showing enrichment in CD41/61^high^ cells versus untreated cells. (C) *SNORD*-inactivated K562 cells (as indicated), were treated with 2nM PMA for 72hrs, and then stained for CD41/61 levels. Left - quantification of all independent repeats of PMA treatment of *SNORD91a*-inactivated cells versus *SNORD98*- or *SNORD127*-inactivated K562 cells. A total of 5 independent repeats were performed. Statistical significance was calculated by Student’s t-test. Right – FACS histograms of a representative experiment showing higher expression of CD41/61 by the *SNORD91a*-inact cells.

To validate these findings, we analyzed our previously generated *SNORD91a*-inact K562 cells alongside *SNORD98*- and *SNORD127*-inact controls, which are not expected to affect megakaryocyte differentiation. Consistent with the screen, *SNORD91a*-inact cells exhibited the highest CD41/61 expression (**Fig.4C**). The increased capacity to differentiate and loss of proliferation upon *SNORD91a* inactivation, was also observed in the HEL erythroid leukemia cell line (**Fig.S3D-E**). Taken together, our findings suggest that Gm4588 regulates cellular state, impacting both proliferation and differentiation.

### 28S-Gm4588 fine-tunes translation to promote stem cell programs

As Gm4588 is positioned within an important site of the ribosome (**Fig.3A**), we next explored its potential role in regulating translation. We found no impact of *SNORD91a*-inactivation on global translation nor on ribosome quantity (**Fig.S4A-B**). As Gm4588 is located in a site related to the binding of elongation factors^78–80^, we performed polysome fractionation and found a lower monosome-to-polysome ratio in *SNORD91a*-inact cells (**Fig.5A**). This might suggest that changes to Gm4588 levels effects elongation rates, leading to frequent initiation events. To explore this further, we performed ribosome profiling of *SNORD91a*-inact cells (**Fig.S4C-D**). Gene set enrichment analysis (GSEA) of RPF, demonstrated positive enrichment for gene sets related to hematopoiesis, translation, mitochondria, and cell cycle in *SNORD91a*-inact cells (**Fig.5B**, and **Supplementary Table 6**). These findings are particularly interesting in the context of HSCs, as HSCs require low protein synthesis rates and low mitochondrial activity for stemness maintenance. Aberrations in these elements lead the cells to exit their stem state and enter the cell cycle^3,8,83–86^. The observed enrichments are also in line with the high responsiveness of *SNORD91a*-inact cells to differentiate upon stimulus (**Fig.3E** and **Fig.4C**). Notably, we found both positive and negative enrichments of signaling-related sets known to be important for cell differentiation (**Fig.5B**), further emphasizing the cellular changes caused by *SNORD91a* inactivation.

**Figure 5.**
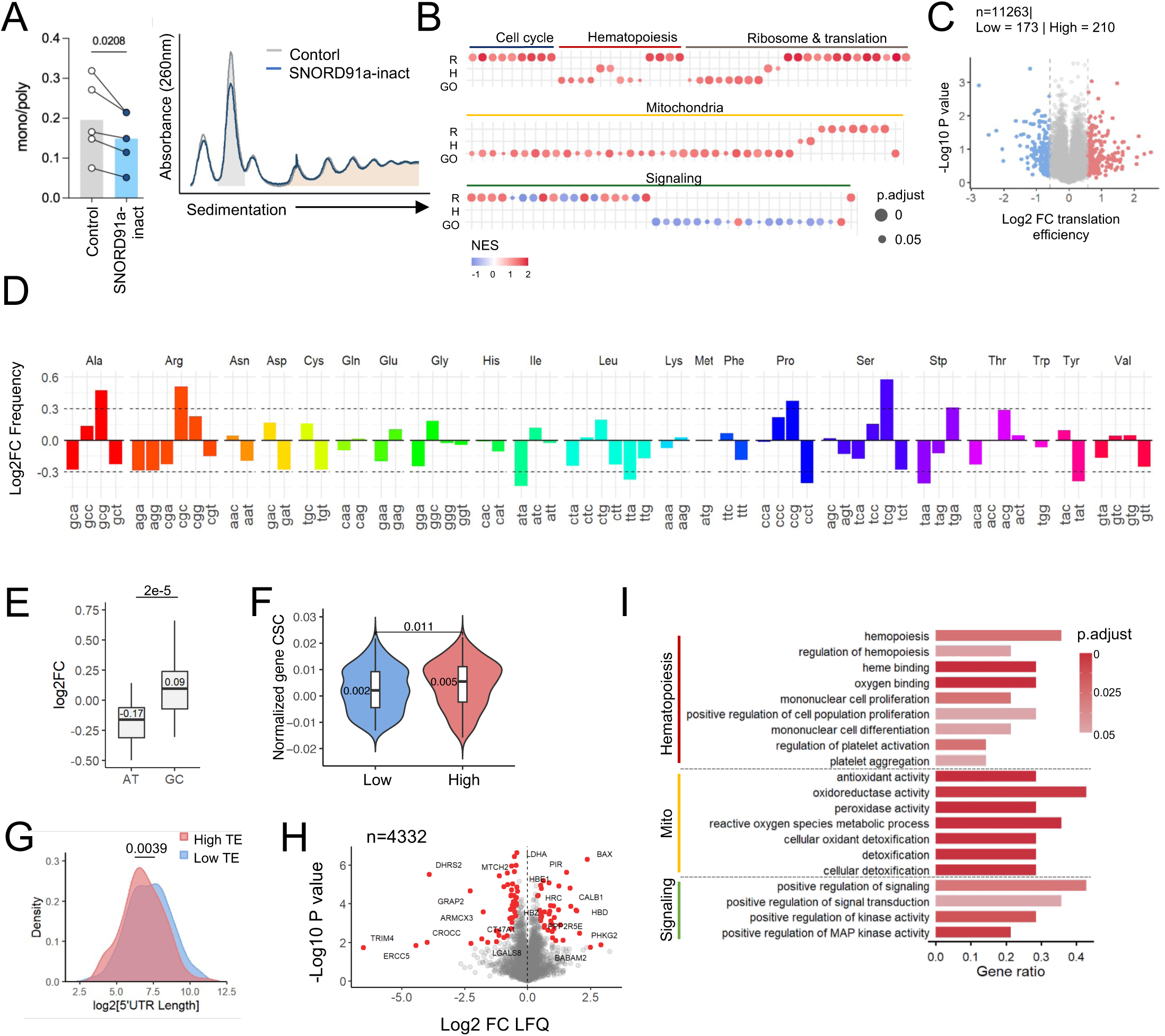
Gm4588 fine-tunes translation to regulate cellular state. (A) Left – quantification of the ratio of the area under the curve of monosomes and polysomes (shaded areas in the left panel). Ratios obtained in the same experiment from the two genotypes are connected with a line (5 independent repeats). Significance was determined by t-test. Right - A representative plot of ribosome fractionation on sucrose gradient. (B) GSEA of RPF in *SNORD91a*-inact cells compared to control. Presented are selected significant gene sets from the Reactome (R), Hallmark (H) and GO databases. Similar gene sets were grouped under a specific category. The full list is in Supplementary Table 6. (C) Differential TE between *SNORD91a*-inact and control K562 cells. Dotted line indicates log2FC of ±0.58 (FC of 1.5). (D) Codon frequency bias of differentially TE transcripts. Transcripts with high TE in *SNORD91a*-inact cells have an increased frequency of G/C-ending codons. (E) Log2 fold change of the frequency of A/T and G/C ending codons. Statistical significance was calculated by t-test. (F) Genes with high differential TE in *SNORD91a*-inact cells are enriched in optimal codons. Comparison of codon optimality score based on Wu et al codon stabilization coefficient (CSC). The score was calculated by summing the codon scores of a gene’s and normalizing to gene length. (G) 5’UTR length distribution analysis of transcripts with differential high or low TE in *SNORD91a*-inact cells (median 184nt in *SNORD91a*-inact versus 247nt in control) Statistical significance was calculated by t-test. (H) Proteomic analysis of *SNORD91a*-inact and control cells. (I) GO analysis of upregulated proteins identified by mass-spectrometry in *SNORD91a*-inact versus control samples. Presented are selected significant GO pathway enriched. Similar pathways were grouped under a specific category. No significant GO pathways were identified using the downregulated proteins.

To gain additional insights into Gm4588-mediated translation regulation, we focused on differential TE (**Fig.5C**). We found a skewing in codon frequency in the high differential TE group in *SNORD91a*-inact cells (**Fig.5D**). Furthermore, we found that these transcripts are enriched in G/C-ending codons (**Fig.5E**), which were previously demonstrated by Wu et al as optimal codons that promote translation specifically in K562 cells^87^. Using Wu et al optimality score, calculated specifically in K562 cells, we identify enrichment of optimal codons in transcripts highly translated in *SNORD91a*-inact cells compared to control (**Fig.5F** and **Supplementary Table 7**). Interestingly, in T-cells, G/C-ending codons were shown to be enriched in differentiating cells compared to proliferating cells^88^. This is in line with the increased differentiation and low proliferation of

### *SNORD91a*-inactivated cells

As the 5’ untranslated region (UTR) bears important elements for translation regulation, we analyzed the differential TE genes and found that transcripts with higher TE in *SNORD91a*- inact cells, have a shorter 5’UTR relative to those with lower TE (**Fig.5G**). We found no differences in the minimum free energy of the 5’UTRs in each group (**Fig.S4E**), suggesting that the differences in TE are not due to secondary structures.

Following ribosome profiling, we performed proteomic analysis which highly correlated with the ribosome profiling data (**Fig.5H** and **Fig.S4F-G**). As in the ribosome profiling data, also in the proteomics analysis pathways related to hematopoiesis, mitochondrial activity and signaling were enriched in *SNORD91a*-inact cells (**Fig.5I**). These findings demonstrate that low Gm4588 levels lead to changes in translational properties, facilitated by codon frequency and 5’UTR length, resulting in an increased differentiation and low proliferation phenotype.

### 28S-Gm4588 regulates mouse HSPCs fitness *ex-vivo* and *in-vivo*

Next, we tested the functional role of Gm4588 in primary mouse hematopoietic stem and progenitor cells (HSPCs), by *ex-vivo* colony formation assay. In contrast to humans, the mouse methylation site is mediated by a single snoRD - *Snord91*. We identified the most efficient sgRNA in the mouse (**Fig.S5A**), and plated transduced HSPCs on semi-solid media (**Fig.6A**). In the first plating we found a mild, yet significant, change in the differentiation of the HSPCs, where *Snord91*-inactivation gave rise to higher proportion of erythrocyte colonies (**Fig.6B**). Importantly, after replating*, Snord91*-inactivated HSPCs produced very few colonies compared to control cells (**Fig.6C**), demonstrating their low self-renewal capacity. In full accordance, CFU assays with *Snord91* over-expressing HSPCs, demonstrated an increase in colony numbers following a significant increase in methylation (**Fig.S5B-C**). These data strongly suggest that high Gm4588 levels, as seen in HSCs (**Fig.2A**), are required for maintaining self-renewal of HSPCs, with a mild impact on their differentiation.

**Figure 6.**
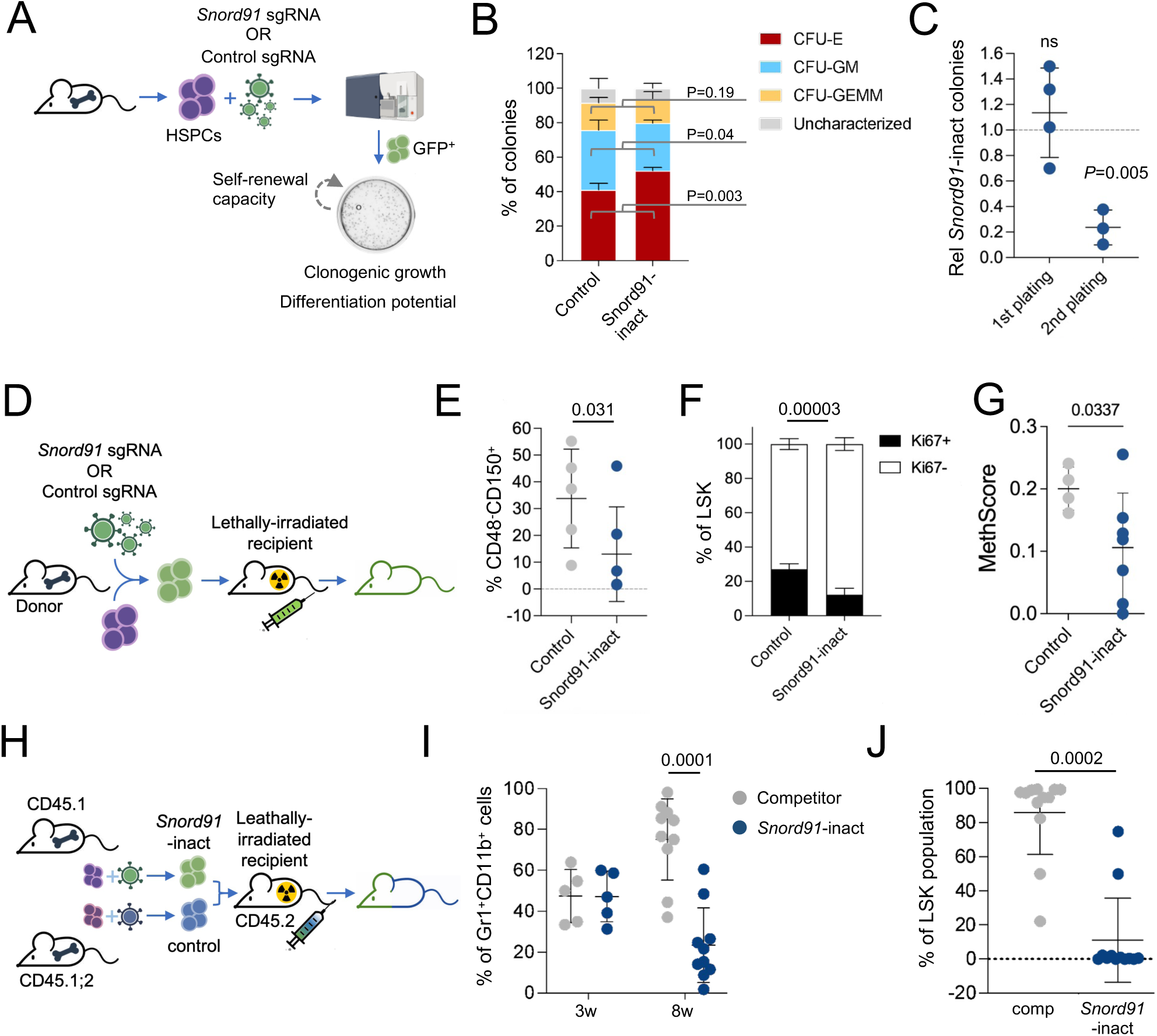
Gm4588/Gm4270 is required for HSC fitness and self-renewal. (A) Schematic depiction of ex-vivo experiment. Isolated BM HSPCs were transduced with a Snord*91*-specific or control CRISPR-Cas9 lentiviral virus. 48hrs after transduction cells were resorted according to GFP expression, and plated in semi-solid media to assess clonogenic growth and differentiation capacity (by morphology). Plates were re-plated to assess self-renewal capacity. (B) Colony characterization in semi-solid media. 10 days after plating colonies were characterized by their morphology. Data are mean±SD of 4 biological replicates, each performed in quadruplets. Significance was calculated by t-test. CFU-E – colony formation unit erythroid, CFU-GM – colony formation unit granulocyte–macrophage, CFU-GEMM – colony formation unit of granulocyte, erythrocyte, monocyte, megakaryocyte. (C) Relative colony number of *SNORD91*- inact HSPCs in the first plating and in the second plating. The number of colonies in the control group was set as 1 and is marked by the dashed line. Data are mean±SD of 4 independent experiments each performed in triplicates. Significance was calculated by t-test. NS – not significant. (D) Illustration of bone marrow transplantation experiments. (E) BM analysis of HSC frequency. (F) Cell cycle analysis, by Ki67 staining of LSK compartment in mice transplanted with Control (n=5) or with *Snord91*-inact (n=7). (G) MethScore of Gm4588 of Lineage-depleted cells extracted from mice transplanted with either *Snord91*-inactivated HSPCs or control cells. Validating low Gm4588 following Snord91-inactivation 4 months post transplantation. (H) Illustration of competitive bone marrow transplantation experiments. (I) Peripheral blood analysis of the differential contribution of each genotype to myeloid cells at 3 weeks post-transplantation (n=5) and 8 weeks post-transplantation (n=10). Each circle represents a single mouse. Bars are mean±SD of all mice in the relevant biological group. Significance was determined by t-test. (J) Bone marrow analysis of competitive transplanted mice at 8 weeks post-transplantation (n=10), showing almost 100% occupancy by the competitor cells. Bars are mean±SD of all mice in biological group. Significance was determined by Student’s t-test. Comp – competitor.

The ability of HSCs to replenish the hematopoietic system is best tested by bone marrow transplantations (BMT). Hence, we performed BMT of single genotype HSPCs, by transplanting either *Snord91*-inact or control HSPCs into lethally irradiated recipients (**Fig.6D**). 2 months after transplantation we found no differences in the major blood lineages (**Fig.S5D**), and no differences in specific cell populations either in peripheral blood or BM (**Fig.S5E-I**). However, at 6 months post-transplantation, when hematopoiesis relays on HSCs rather than HSPCs, BM analysis of the transplanted mice demonstrated reduced numbers of HSCs (**Fig.6E**), with higher percentage of cells at G0 (Ki67-negative, **Fig.6F**). At this time point we extracted RNA from Lineage-depleted BM cells and validated low Gm4588 methylation in the *Snord91*-inact transplanted mice (**Fig.6G**).

To further test HSC fitness, we performed competitive-BMT (**Fig.6H**). Three weeks after transplantation peripheral blood cell populations and found that *Snord91*-inact HSPCs contributed 50% of the cells in the myeloid compartment (**Fig.6I**). However, at 8 weeks post-transplantation, we found very little contribution by *Snord91*-inact HSPCs (**Fig.6I**). In full accordance, BM analysis at 8 weeks showed that the *Snord91*-inact cells were outcompeted, as we found almost 100% occupancy of the BM by the competitor (**Fig.6J**). Thus, we conclude that while *Snord91*-inact HSPCs can facilitate short-term reconstitution, these cells have very low fitness and are easily outcompeted. This demonstrates that Gm4588 is required in high levels to maintain a stem gene-expression and is essential for HSC fitness.

### Dynamics of Gm4588 beyond the erythroid compartment

As in HSCs, also in other stem cells, protein synthesis is restricted in the stem cell population and increases upon differentiation^8,89–91^. Thus, our data offers two possible models for rRNA methylation in stem cells – the first, that there are common stem-methylations assigned to regulating stem cells’ translation. And the second, that each type of stem cell has its unique methylation profile to tailor its translational output.

To assess if low levels of Gm4588 are specific to erythroid differentiation or rather is a general phenomenon in the hematopoietic system, we sorted specific populations of B-cell development which occurs within the BM. We sorted HSCs, CLP (common lymphoid progenitors), and preB-cells (committed B-cells precursors). Using RiboMethSeq, we found that, similar to erythrocyte differentiation, B-cell differentiation also exhibits variable methylation sites (**Supplemental Table 5**). Moreover, we found that these sites, including Gm4588, show a decrease in methylation during differentiation (**Fig.7A-B**).

**Figure 7.**
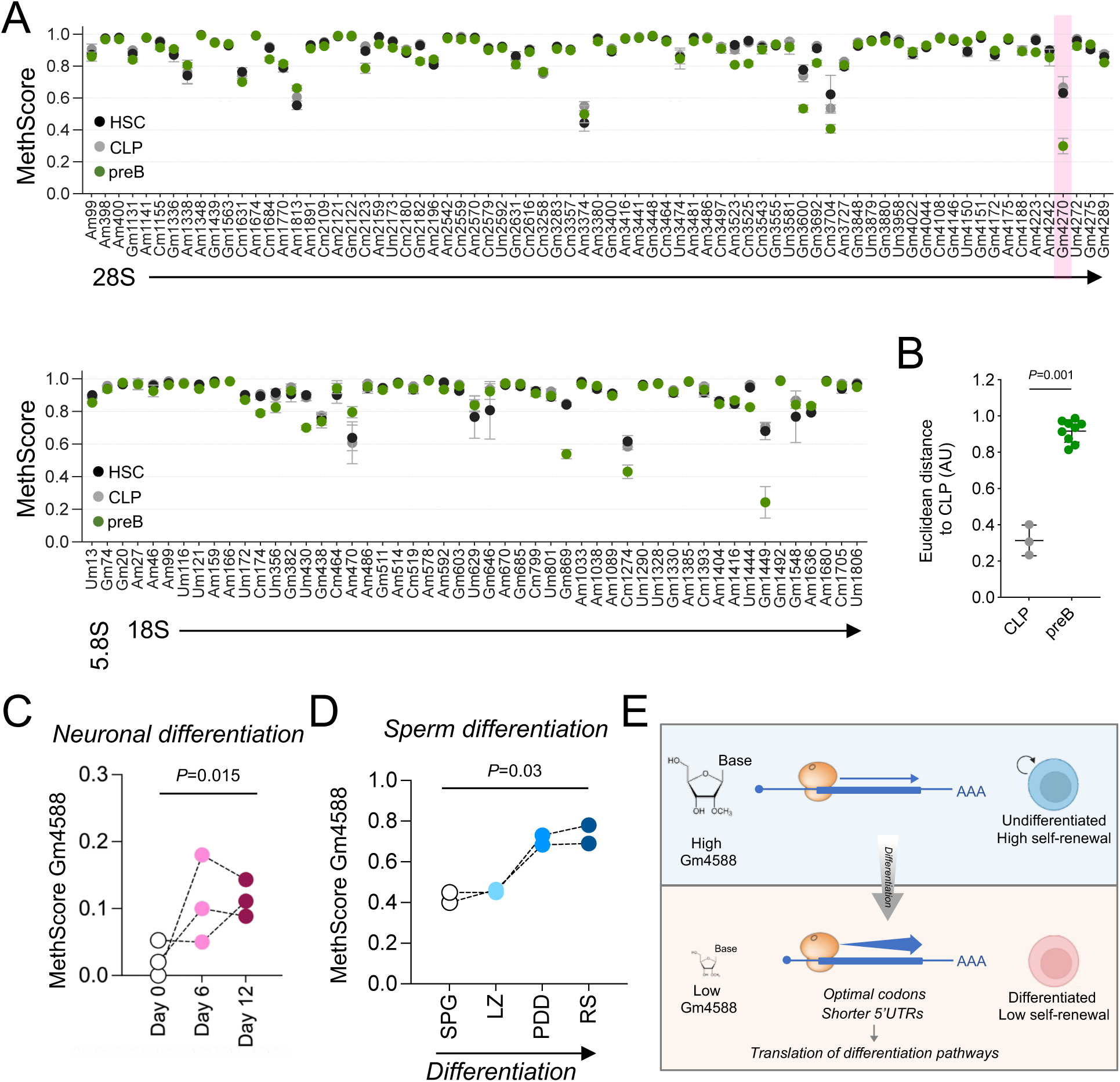
Loss of Gm4588 methylation levels defines hematopoietic differentiation. (A) RiboMethSeq analysis of B-cell precursors. Total RNA was extracted from HSC (Lin^-^ Sca1^+^cKit^+^CD48^-^CD150), common lymphoid progenitors (CLP, Lin^-^Flk2^+^IL7R^+^) and preB-cells (Gr1^-^ CD11b^-^TER119^-^CD3^-^B220^+^CD43^-^IgD^-^IgM^+^) which were sorted from the bone marrow of 8-10 weeks old mice. Shown are mean±SD of three independent experiment, each replicate consisted of pooled 8 mice. (B) Euclidean distance analysis to quantitively evaluated dissimilarities of 2’OMe profiles. Significance was determined by t-test relative to CLP. (C) Gm4588 levels in mESC undergoing cortical differentiation. Total RNA was extracted at Day 0, Day 6 (induction phase) and day 12 (neurogenesis stage) and analyzed by RiboMethSeq. Three biological replicates were performed. (D) Gm4588 levels in Spermatogonia (SPG, 2N and 4N), leptotene/zygotene (LZ, 4N), pachytene/diplotene/diakinesis (PDD, 4N), and round spermatids (RS, 1N) representing 4 staged of sperm differentiation from 2N stage, through 4N and ending at 1N. Shown is the methscore of two biological replicates, each experiment consisted of 3 mice. (E) A model for the translational modulation by Gm4588. In undifferentiated cells there is high methylation levels of Gm4588. During differentiation, methylation of Gm4588 is lost, supporting the translation of differentiation-related transcripts and leading to loss of self-renewal capacity.

We next examined additional stem cell types and differentiation pathways beyond the hematopoietic system. Specifically, we evaluated Gm4588 levels during differentiation of mouse embryonic stem cells (mESCs) into cortical neurons^92,93^ and during spermatogenesis, where cell-type–specific ribosomes have been described^94^. We found that unlike in the hematopoietic system, Gm4588 is completely unmethylated in mESCs and present at very low levels in spermatogonia (**Fig.7C-D**). Interestingly, in both neuronal and sperm differentiation, Gm4588 methylation is dynamic, but in contrast to the hematopoietic system, it increases during these processes (**Fig.7C-D**). Together, these findings support the second model and suggest that the 2ʹOMe landscape is distinctly regulated across stem cell populations to fine-tune and tailor translation.

## DISCUSSION

Here we analyzed the transcriptome, translatome, and rRNA methylome of rare populations along erythroid differentiation, using ultra-low ribosome profiling and comprehensive mapping of 2’OMe landscape. Our results confirm previous reports of specific translational capacities in HSCs and progenitors^8^ (**Fig.1H**), and provide higher resolution into the regulation of individual transcripts. Additionally, we show that changes in translational capacity can be cell-type-specific and serve to distinguish discrete cell types (**Fig.1K**). We propose that dynamic regulation of 2ʹOMe contributes to this cell-type specificity. By linking 2ʹOMe to cellular function, we identify methylation at Gm4588 as essential for maintaining stem cell-related gene expression and fitness *in-vivo*. Mechanistically, we found that low Gm4588 methylation promotes translation of differentiation-associated transcripts, rich in G/C-ending codons^87,94^ and with short 5’UTRs. Ultimately resulting in loss of proliferation and increased cellular differentiation (**Fig.7E** and **Fig.5B**)

Previously, Zhou et al^95^, have identified a set of 2’OMe sites which are highly methylated in leukemia stem cells, among which were 28S-Gm4588 and 18S-1447. Here we identified these two sites as very dynamic, with high methylation levels in HSC, that are significantly lost during differentiation (**Fig.2A**). We suggest that our findings reveal the mechanism which was hijacked by leukemia stem cells – high 2’OMe is required for maintenance of stemness. We propose that the leukemic cells mimic the HSCs’ 2’OMe rRNA profile to maintain their stem gene-expression and to sustain malignancy. Otherwise, if high methylation levels are not maintained, self-renewal capacity will be lost, as we have demonstrated by inactivation (**Fig.6H-J**), and by over-expression experiments (**Fig.S5B**). Our finding, that non-hematological stem cells also differentially regulate specific methylation sites (**Fig.7B-C**) is in line with recent work showing that rRNA modification patterns can be used for tissue identification, and discrimination between normal and malignant cells^96^. Our study together with others ^10,70,96^, support rRNA modification profiles as an element tailored to support cell identity across various biological systems.

We establish a direct role for a specific methylation site in HSC fitness, by genetically altering the corresponding snoRNA. Hypothetically such alterations can also be due to mutations in the snoRNA sequence itself or in the sequence elements required for snoRNA maturation.

Given the importance of rRNA methylation to HSCs, this suggests that snoRNAs are potential candidate genes for underlying human bone marrow failure disorders, such as ribosomopathies, as their loss-of-function might result in aberrant hematopoiesis. Interestingly, a recent publication has used whole genome sequences of patients with neurodevelopmental disorders to identify a mutation in a non-coding RNA, a small nuclear RNA, as a prevalent mutation and link it to neurodevelopmental disorders^97^. This further underlines the potential widespread role of non-coding genes in human pathologies and the need for their mapping and investigation.

It is interesting to consider that changes to the ribosome itself by regulation of its rRNA modifications profile (and hence changes to translation), represent a novel facet of decision making - where specific changes to rRNA modifications drive one type of cell fate choice, while other changes drive the other, like we have demonstrated with MkP (**Fig.1** and **Fig.2**). In this respect, while we have identified changes in translational capacity in MkP relative to HSC (**Fig.1H- K**), we have not found dynamics of 2’OMe (**Fig.2**). This might be due to changes to other rRNA modifications, such as pseudouridine, which are also abundant on rRNA and regulate translation^19,98^. Such complementary mapping, of additional rRNA modifications, will provide significant insights into translation-based regulation of cell identity.

## Supporting information

Supplementary Table 1 - ULrna_seq_tpm_counts

Supplementary Table 2 - ULribo_seq_tpm_counts

Supplementary Table 3 - ULriboseq RPF and RNAseq deseq top 50 signature genes

Supplementary Table 4 - top and bottom 5 precent TE genes

Supplementary Table 5 - P-adj values of all RMS

Supplementary Table 6 - K562 ribo seq GSEA full results table

Supplementary Table 7 - Optimality score of K562 ribo seq 210925

## ACKNOWLEDGEMENTS

DN is supported by grants from the Israel Science Foundation personal research, Grant No. 1193/22, and European Research Council, grant no. 10107560. DN Received support from the Abisch-Frenkel Faculty Development Lectureship and by the Alon Fellowship from the Israeli Council for Higher Education. DN is a member of the Human RNome Consortium. We acknowledge the use of BioRender.com for creating illustrations. We thank Michal Rabani for her assistance and advice in bioinformatic analyses.

## AUTHOR CONTRIBUTIONS

O.R., S.B.D, J.W, M.L. and D.N., designed and performed research. M.A. H.A., N.R., N.U., A.R., performed experiments. V.M., and Y.M., performed mouse hematopoietic RiboMethSeq and analysis. G.M., S.S., C.P., and C.M contributed vital reagents. N.M. and O.K provided vital assistance. W.B.M, M.V., and A.O ultra-low ribo-seq, sample preparation and analysis. O.R., S.B.D, Y.H. and Y.L. performed vital data analysis. D.N. wrote the manuscript.

---

**Supplemental Figure 1.**
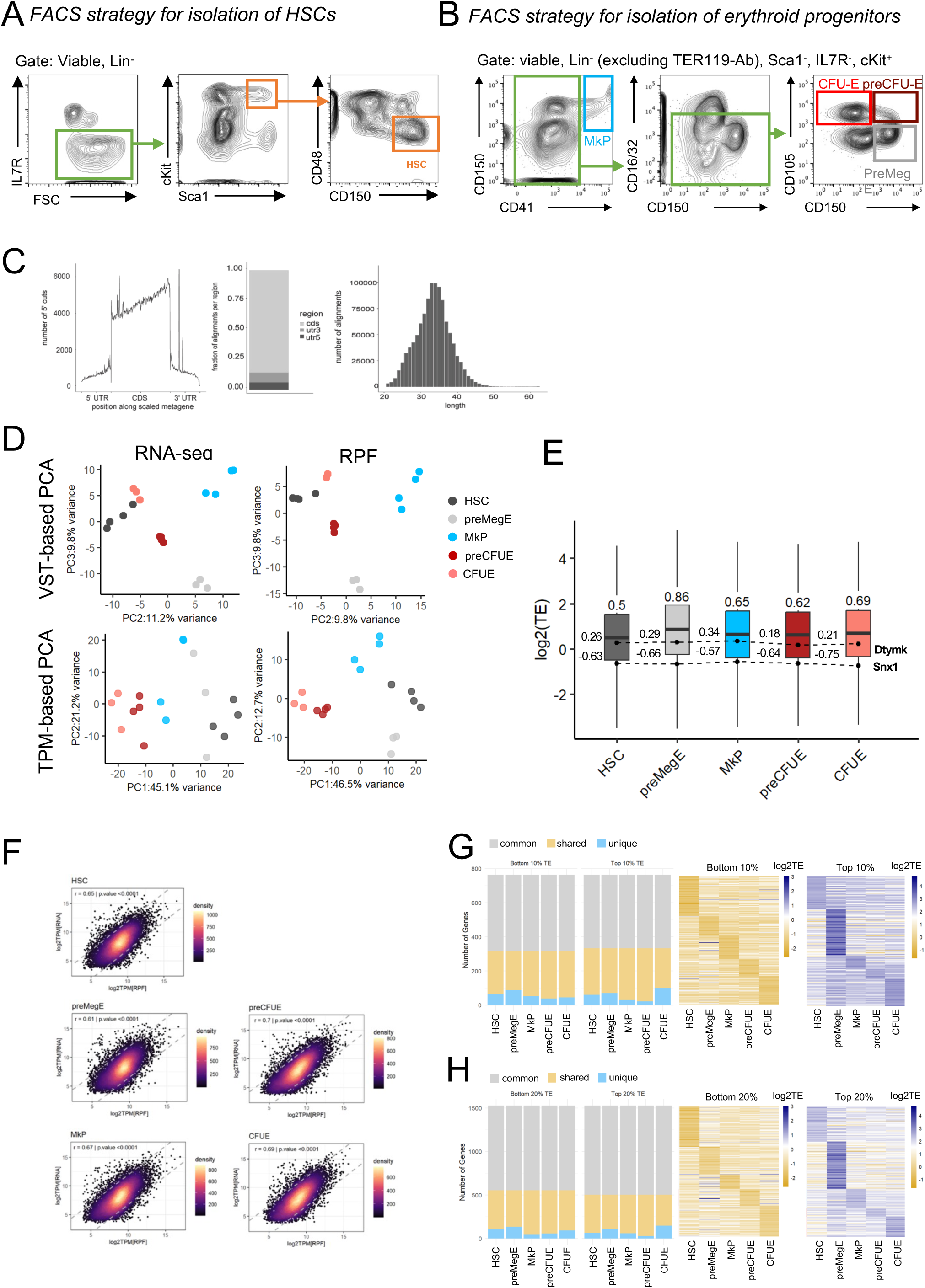
Ultra-low ribosome profiling. (A and B) FACS gating strategy for sorting HSCs (A) and erythroid progenitors (B). (C) Quality control assessment of a representative sample of ultra-low RPF sequencing (ribo-seq). left and middle - read alignment on the transcript (5’UTR, CDS, 3’UTR), right – length distribution of aligned reads. (D) PCA of ultra-low RNAseq (left) and ultra-low ribo-seq (right). Top - shown are the plots of PC2 vs PC3 of DESeq variance stabilizing transformed (VST) reads. Bottom – PCA using TPM values showing superior clustering using RPF data (ribo-seq). (E) Global TE in specific cell types, with marks for normalizing housekeeping genes that their TE does not change amongst the cell types. No significant differences in the TE of the normalizing genes (Dtymk: p = 0.4, Snx1: p = 0.4, Kruskal–Wallis test). (F) Plots of RFP TPM vs RNA-seq TPM showing 5% of genes for each cell type indicated on top. (G) Left panel - Top and bottom 10% of TE genes, 3 categories – genes which are shared by all cell types (gray), that are shared by several but not all cell types (gold), and those which are uniquely in the top/bottom of a single cell type (blue). Heat map of the unique TE genes of the top (middle) and bottom (right) 10% according to cell type. (H) same as G for the top and bottom 20% of TE genes.

**Supplemental Figure 2.**
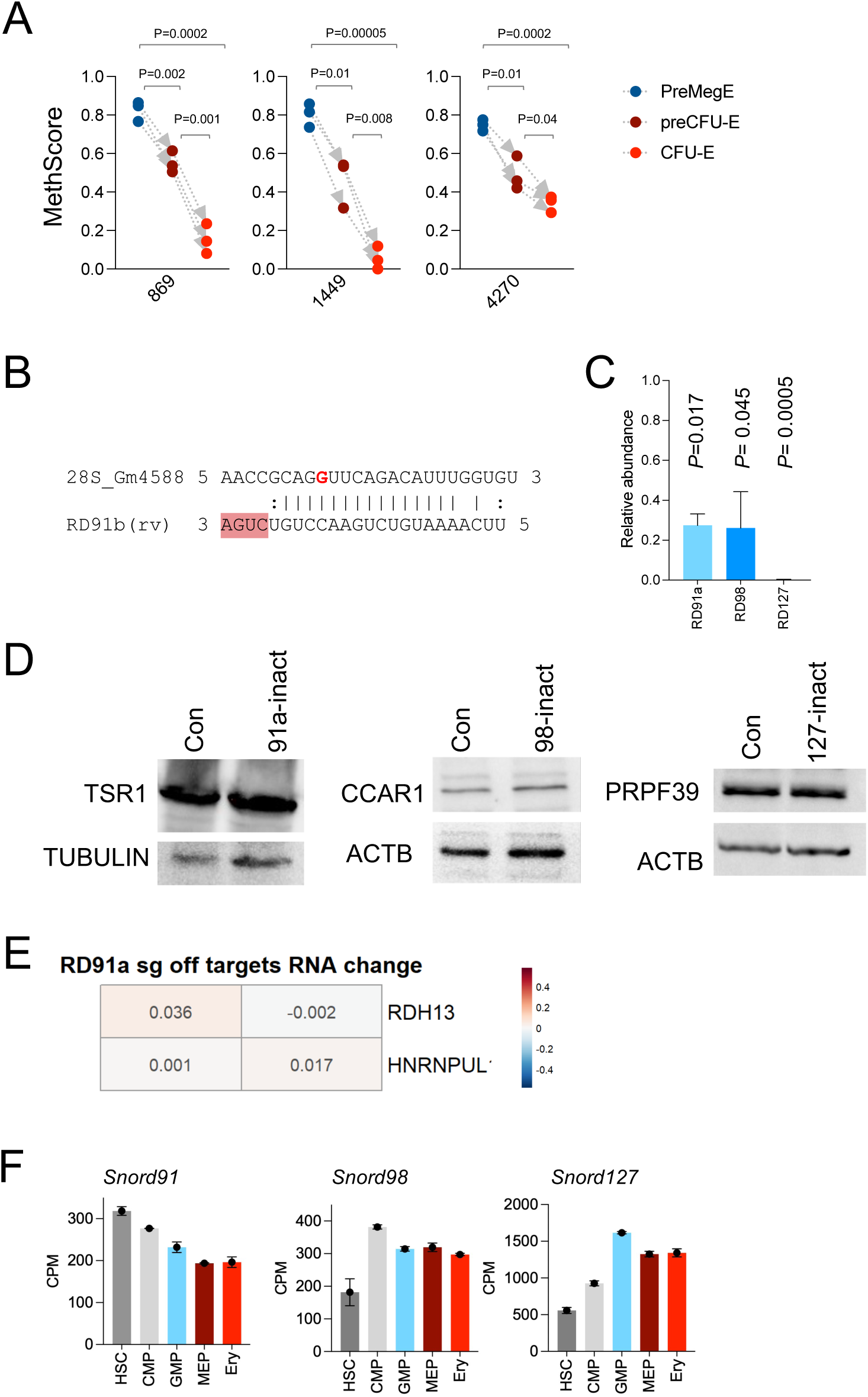
Generation of methylation-specific inactivated cells. (A) MethScore of 18S-Gm869, 18S-Gm1449 and 28S-Gm4270 (mouse nomenclature) in the indicated hematopoietic compartments. Significance was determined by t-test. (B) Alignment between *SNORD91b* and its corresponding methylation site 28S-Gm4588. Highlighted in dark pink is the snoRNA’s D’ box. In red is the modified nucleotide on rRNA. (C) qPCR of the relevant snoRNA’s abundance in the indicated SNORD-inactivated K562 cells. Significance was determined by t-test. (D) Western blot analysis of the corresponding host gene in each SNORD-inactivated cell. (E) Fold change (RNA-seq) of the 2 off targets predicted for the sgRNA targeting SNORD91a (by Benchling). No significant change to RNA levels were observed. (F) CPM values of the indicated Snord in various hematopoietic compartments. Data reanalyzed from *Wang et al, Nature Cell Biology, 2025*.

**Supplemental Figure 3.**
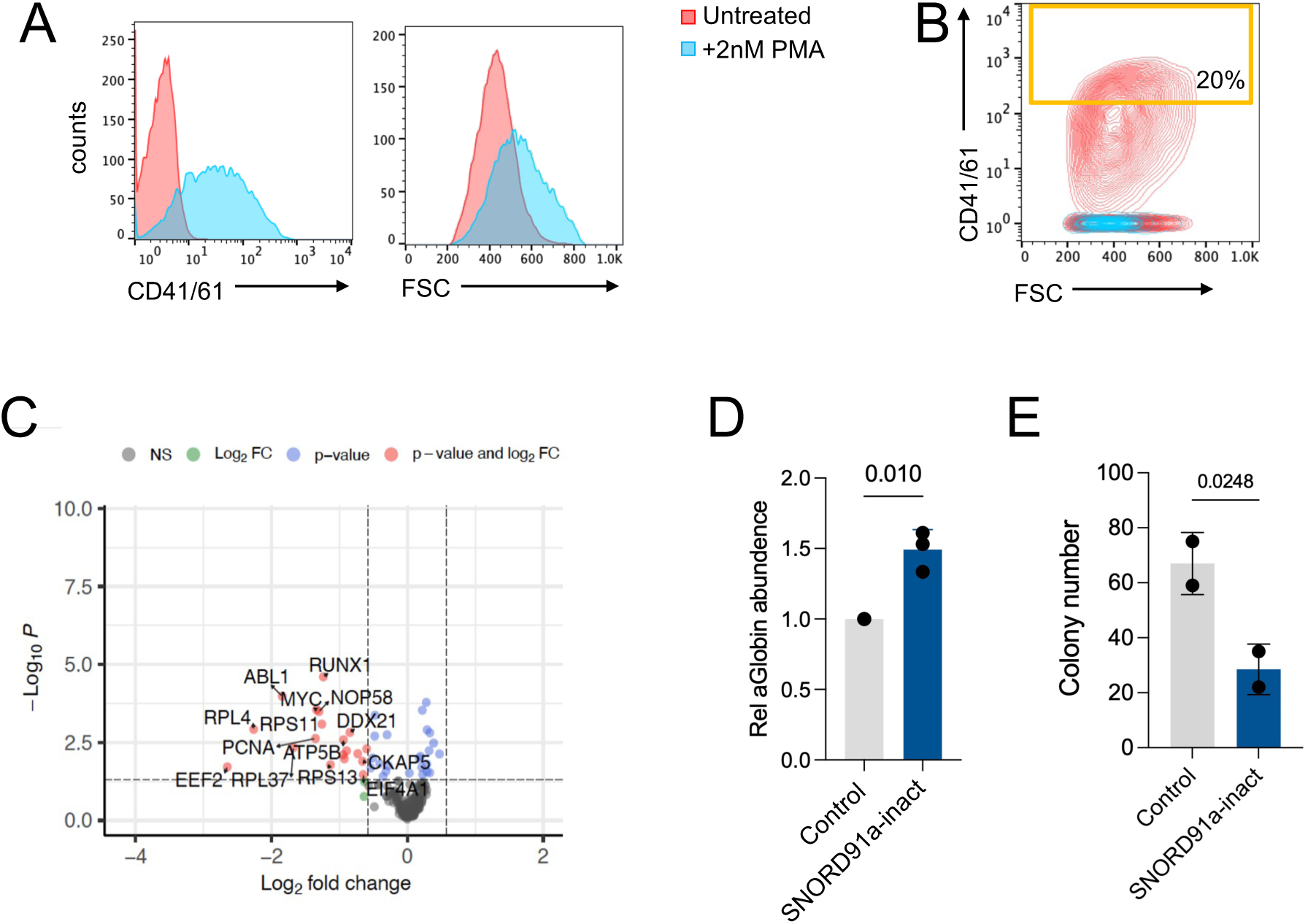
Human SNORD CRISPR screen. (A) FACS analysis of K562 cells treated with 2nM PMA for 72hrs showing an increase in CD41/61 expression (left) and in cell size (right). (B) FACS plot of one of the biological replicates of CD41/61 staining of library-transduced K562 cells treated with PMA. Rectangle represents the sorted gate of CD41/61^high^ cells. (C) Volcano plot of protein-coding genes which were significant in the SNORD CRISPR screen. (D) levels of alpha-globin in hemin-treated HEL cells (erythroid leukemia cell line) either transduced with SNORD91a sgRNA or a control sgRNA. Three independent biological replicates were performed. (E) Colony formation assay of HEL cells (erythroid leukemia cell line) either transduced with SNORD91a sgRNA or a control sgRNA. Two independent biological replicates were performed.

**Supplemental Figure 4.**
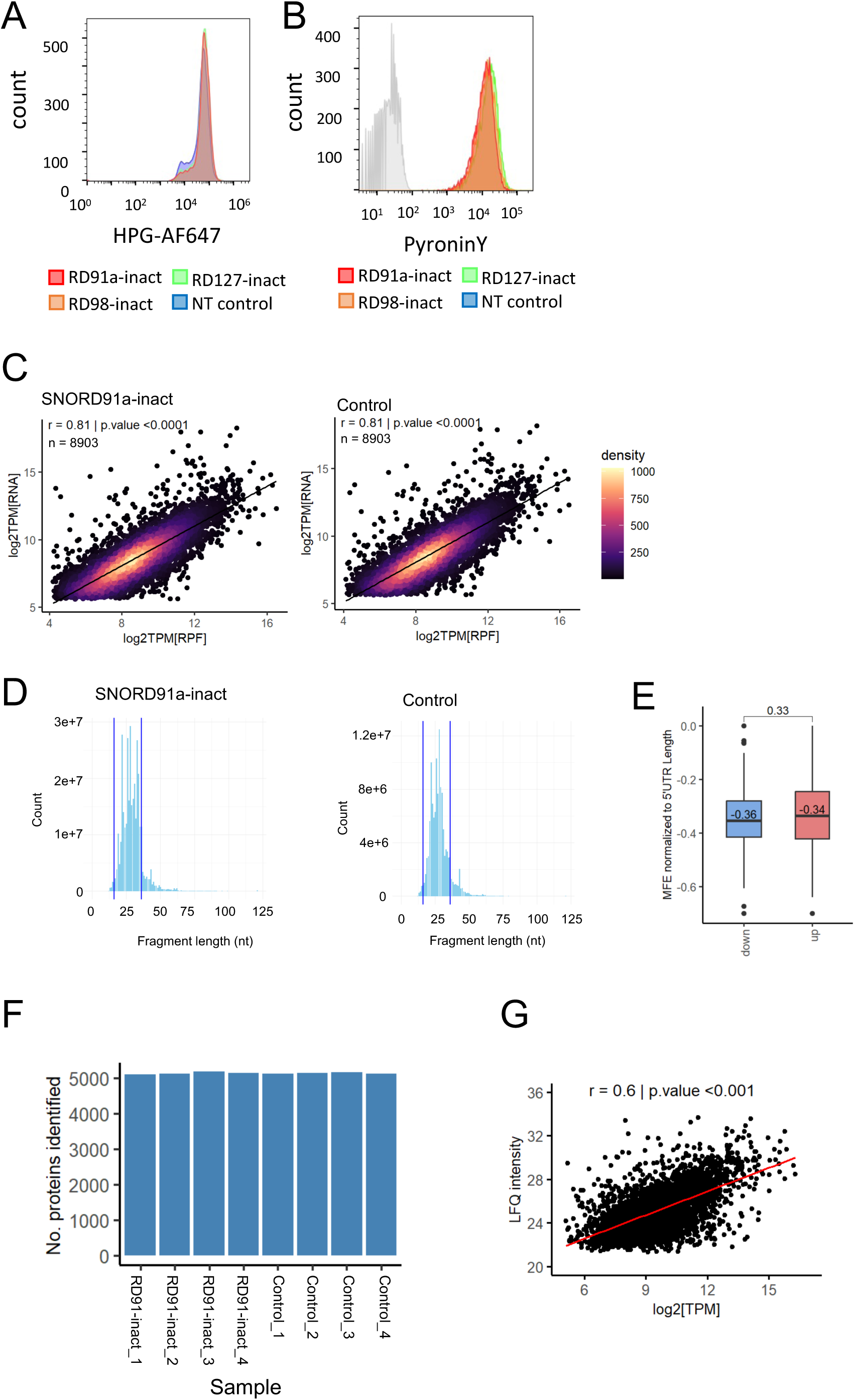
Translational analysis of Gm4588-low K562 cells. (A) Representative FACS histogram of HPG pulse in the indicated SNORD-inactivated cells. After treatment cells were processed and clicked to Alexa-Flour 647 and analyzed by FACS. Data shows no difference between the inactivated cells. (B) Cellular RNA content. Cells were stained with PyroninY for total RNA analysis (reflective on ribosome numbers). Cells were first stained with Hoechst33324 to block binding of PyroninY to DNA, then PyroninY was added and analyzed by FACS. (C) Correlation between RNA-seq and RPF of SNORD91a-inactivated (left) and control (right) cells. (D) Quality control of ribosome protected fragments of K562 cells, as indicated on top. Read length distribution is primarily within expected range (16-36 b, blue lines). (E) Minimum free energy of predicted 5’UTR secondary structures (ViennaRNA Package) of transcripts with differential high or low TE in *SNORD91a*-inact cells, show no significant difference. Statistical significance was calculated by t-test. (F) Number of proteins identified in proteomics analysis (Razor+unique > 0). Similar number proteins were identified in all samples. (G) Correlation between RPF and proteomics of SNORD91a-inactivated cells.

**Supplemental Figure 5.**
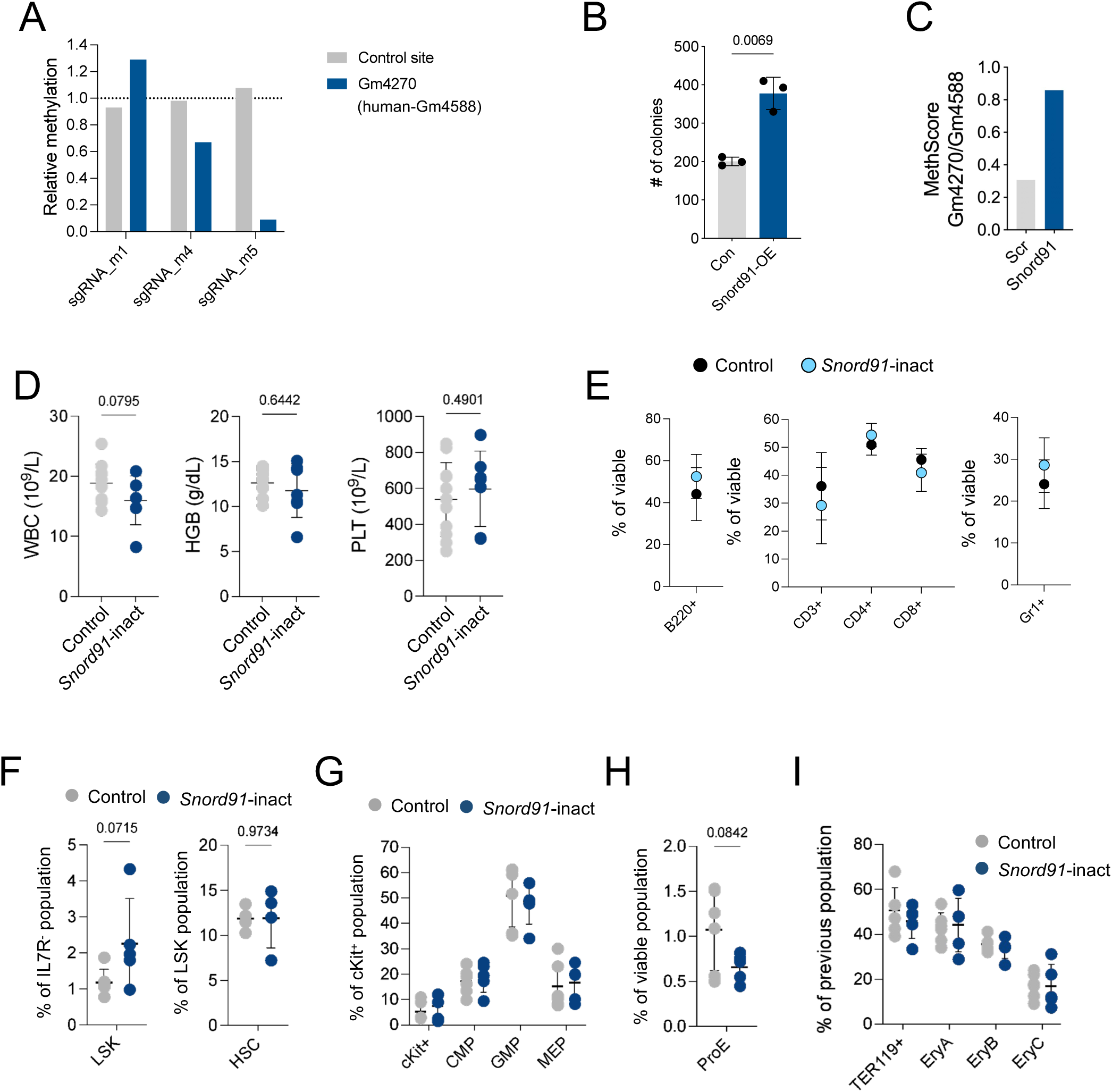
S*n*ord91-inactivated HSPCs facilitate short-term reconstitution of the bone marrow. (A) Relative 2’OMe levels, assessed by RTL-P of mouse embryonic fibroblasts transduced with various sgRNAs targeting mouse-*Snord91*. (B) ex-vivo colony formation assay of HSPCs overexpressing either *Snord91* or a scrambled sequence of *Snord91* as control. (C) MethScore of Gm4270 (Gm4588 in human nomenclature) of LSK cells transduced either with *Snord91*-scramble or *Snord91*, a week after plating in Methocult for CFU assay. (D) Complete blood count analysis of control and *Snord91*-inact transplanted mice 8 weeks post-transplantation. Each circle represents a single mouse. White blood count – WBC, red blood cells – RBC, hemoglobin – HGB, platelet – PLT. No significant differences were found. (E) Peripheral blood FACS analysis of bone marrow-transplanted mice, at 8 weeks post-transplantation. Data are mean±SD of 4-6 mice. (F-I) Bone marrow analysis of bone marrow-transplanted mice, at 8 weeks post-transplantation. Each circle is a mouse. No significant differences were found in all analyses.

## METHODS

### Ultra-low ribosome profiling

Ultra-low ribosome profiling was performed by FACS sorting 1000 cells directly into cooled lysis buffer (Tris pH 7.5, MgCl2, CaCl2, NaCl, Triton X-100, RNase IN plus, cycloheximide). Cells were flash frozen and sorted at -80. Frozen samples were shipped to Single Cell Core (An Oncode Institute Facility) for further processing and analysis. Ribosome profiling libraries were prepared using a version of the scRibo-seq^1^ protocol that was adapted for higher cell inputs (1000 cells). To achieve this, we scaled up all reaction volumes 100×. To process the libraries, cells were first FACS sorted into 0.5 mL DNA LoBind tubes containing 4.5 µL of lysis buffer (22 mM Tris pH 7.5, 16.5 mM MgCl2, 5.5 mM CaCl2, 165 mM NaCl, 1.1 % Triton X-100, 2.2 U/µL RNasein plus (Promega), 0.11 mg/mL Cycloheximide). Then, 5 µL of Micrococcal Nuclease (MNase, 62.5 U/ µL, New England Biolabs) and RNasein Plus (1.2 U/µL RNasein Plus) was added to digest the exposed RNA. The remaining steps of the protocol (proteinase K, end repair, 3ʹ ligation, 5ʹ ligation, reverse transcription, PCR, size enrichment, and purification of libraries) were performed as described in scRibo-seq^1^. The concentrations of the pre-adenylated 3ʹ adapter (OMV630_miRNA4_3App) and 5ʹ adapter (OMV632_miRNA5_10U) were increased 2× in the 3ʹ ligation mix and 5ʹ mix, respectively, to adapt to the higher number of cells. Ribo-seq libraries were sequenced using Illumina NextSeq 2000 with 72 cycles for read 1, 6 cycles for the i7 index read, and 10 cycles for the i5 index read (sample index).

Ultra-low RNA seq was performed by sorting 10,000 cells, from the same sample as the ultra-low ribosome protected fragments, into lysis buffer. RNA extraction was done with the RNA XS NucleoSpin (by MN). Prior to library prep RNA integrity was measured using Pico Bioanalyzer, only samples with a RIN >8 were further processed. Library prep was performed using SMARTseq LP (Katarabio). Samples were sequenced on NovaSeq.

### Assessing 2’OMe levels (RiboMethSeq and RTL-P)

0.2-2 ug total RNA was fragmented by alkaline hydrolysis in a final concentration of 50mM bicarbonate, and heated to 95C for 10 minutes. To stop the fragmentation 1ml of cooled participation mix (1ml absolute ethanol, 1% 3M NaOAc, and 1% glycogen per reaction) was added, and samples were flash frozen in liquid nitrogen. For precipitation, the mix was centrifuged for 30 min at 4C and subsequently washed with 80% ethanol. For RiboMethSeq of mouse progenitor cells, 4 mice were pooled for each biological repeat and cells were FACS isolate. RNA was extracted using the RNeasy kit (Qiagen). End repair was performed using Antarctic phosphatase and T4 PNK prior to library preparation by NEBNext small RNA library kit. RTL-P was performed as previously described in ^2^. Final dNTPs concentration was 500uM for high concentration reaction and 0.5uM for low concentration reactions.

### Cell culture, differentiation and clonogenic growth assays

K562 cells (ATCC) were grown in RPMI (Gibco) with 10% FBS, 5% penicillin-streptomycin, 5% L-Glutamine Solution (Sartorius), 5% Sodium Pyruvate Solution (Sartorius) and 5% MEM EAGLE (Sartorius). Cells were kept in 37 °C and passaged 1:10 twice a week. For differentiation assays cells were cultured with 30µM Hemin (Sigma-Aldrich) or with 2nM PMA (Sigma-Aldrich) for 72h to induce erythroid or megakaryocyte differentiation, respectively. Cells were collected and RNA was extracted for qPCR to assess gamma-globin levels (hemin-induced differentiation), or were stained for megakaryocyte markers CD41/61 (clone A2A9/6) in flow cytometry (PMA-inducted differentiation). In colony assay 500 cells were plated in 1.1ml MethoCult H4034 plated (Stem cell technologies)

### Plasmids transfection and lentivirus

SgRNAs targeting the human or mouse snoRNAs were designed using the Benchling CRISPR design tool. The gRNAs were cloned into lentiCRISPRv2-puro (a gift from Brett Stringer^3^, Addgene plasmid #98290) or lentiCRISPRv2GFP (a gift from David Feldser^4^, Addgene plasmid #82416). Plasmids were transfected into HEK293T cells along with packaging plasmids psPAX2 (a gift from Didier Trono, Addgene plasmid #12260) and pMD2.G (a gift from Didier Trono, Addgene plasmid #12259) using LT-1 or PEI-MAX transduction reagents (PEI MAX;Clini Sciences). Lentivirus particles were harvested after 48 and 72 hours. For mouse sgRNA, lentivirus particles were concentrated by ultracentrifugation using SW28 rotor, conical tubes with the proper adapters, at 25,000 RPM for 2 hours, and resuspended in StemSpan media (Stem cell technologies).

### Mouse HSPCs isolation and transplantation

Long bones from 8-week-old mice were collected and crushed. The cells were treated with ACK, washed with PBS+2%FBS and then subjected to further analysis. Lineage^+^ cells were depleted by mouse lineage depletion kit (Miltenyi Biotec) and then sorted on an AriaIII BD sorter using antibodies mentioned in *Antibodies for flow cytometry*. LSK cells were then transduced with ultracentrifuge-concentrated lentiviruses, by centrifugation at 37deg for 30min, and incubated in full StemSpan media (supplemented with IL-3, IL-6, TPO and SCF). 48hrs after transduction cells were sorted according to GFP positivity. For colony formation unit (CFU) assays a total of 800 GFP^+^ cells were plated in 1ml MethoCult M3434 optimum medium (stem cell technologies) in 35 mm Petri dishes in quadruplicates. Plates were incubated for 7 days. To visualize the colonies, MTT reagent (Thermo Fisher Scientific) was added to three replicates for 1 h. Colonies were scored using a Epson scanner and ImageJ’s cell counter. The fourth plate was used for replating 10,000 cells.

For bone marrow transplantations 8–10-week-old C57BL/6J (CD45.2) recipient mice were pre-treated with baytril for two days prior to lethal irradiation by two doses of 550 cGy (with a 4 hours interval) using an XRAD320 irradiator. Mice were injected to the tail vein with a total of 10,000 GFP^+^ LSK cells in addition of 300,000 supporting BM cells (including RBCs).

### Antibodies for flow cytometry

We used monoclonal antibodies from Biolegend specific for the following: ScaI (D7); c- Kit (2B8); CD150 (TC15-12F12.2); CD135 (Flk2; A2F10); CD4 (RM4-5); CD16/32 (93); CD34 (RAM34); CD71 (R17217); TER119 (Ter-119); CD3e (145-2c11); CD11b (M1/70.15.11.5); B220 (RA3-6B2); Gr-1 (RB6-8C5); CD45.1(A20); CD45.2(104); IL7R (A7R34); CD43 (S11); IgD (11-26c.2a) ; IgM (R6-60.2); Ki-67 (16A8) and Zombie Aqua for cell viability. For lineage detection, we used Hematopoietic Lineage Labeling Cocktail conjugated to Biotin (Miltenyi Biotec, 130-092-613), and either SAV-APC-cy7 or SAV-PE-cy5. For intracellular staining we used BD Cytofix/Cytoperm™ Plus Fixation/Permeabilization Solution Kit with BD GolgiPlug™ (555028) according to manufacturers instructions.

### Isolation of mouse male germ cells

The isolation of male germ cells was performed exactly as detailed in ^5^. In summary, hemizygous Stra-iCre males were mated with homozygous tdTomato females, and 3- to 4-month-old males positive for Stra-iCre were utilized as the source of testis tissue. In these mice, tdTomato expression was confined to germ cells. Due to the decreasing intensity of tdTomato expression during spermatogenesis, distinct germ cell populations could be isolated using flow cytometry sorting. Seminiferous tubules were collected from six mice and subjected to enzymatic digestion with collagenase and trypsin. After removing dead cells with the Dead Cells Removal Kit (130- 090-101, Miltenyi), four populations of germ cells at different stages of differentiation were obtained: spermatogonia (SPG), leptotene/zygotene (LZ), pachytene/diplotene/diakinesis (PDD), and round spermatids (RS). The SPG population consists of diploid undifferentiated spermatogonia, as well as differentiating spermatogonia that replicate their DNA without undergoing division. The LZ and PDD cells are tetraploid and represent sequential stages of prophase of meiosis I. Finally, RS cells are haploid and result from two rounds of reductive meiotic division.

### Culture of mESC

The culture and differentiation of mouse embryonic stem cells (mESCs) were conducted as previously described ^6^. Briefly, the E14-Tg2A mESC line was cultured on plates coated with 0.1% gelatin in ESC medium. Cells were maintained in a feeder-free culture system using 2i medium, which consisted of ESC medium supplemented with 1 μM PD0325901 and 3 μM CHIR99021 (both from Peprotech). Differentiation into cortical progenitors and neurons was achieved through an in vitro corticogenesis protocol following the method described by Gaspard et al^7^. For RNA extraction, cells were collected at three differentiation stages: before differentiation (D0), after six days (D6), and after 12 days (D12). According to Gaspard et al^7^, D6 corresponds to the neural induction period, while D12 represents a neurogenesis stage. Total RNA was isolated using RNeasy Mini Kits (Qiagen).

### Pyronin Y assay RNA quantification

Pyronin Y staining was performed as follows: cells at ∼50% confluence were stained with 5 μg/ml of Hoechst 33342 (Sigma-Aldrich) in Hoechst staining buffer (Hank’s Balanced Salt Solution + 3% FCS + 10 mM HEPES) at 37 °C for 45 min. Then pyronin Y (0.5 μg/ml; Sigma-Aldrich) was added to the buffer and the cells were further incubated for 45 min. cells were then collected for FACS analysis.

### Polysome fractionation and standard ribosome profiling

Cells were then incubated with cycloheximide (CHX) at a final concentration of 100 µg/ml for 15 min, then washed with ice-cold PBS containing 100 µg /ml CHX. Cells were pelleted by centrifugation and subsequently lysed in polysome lysis buffer (20 mM Tris-HCl, pH 7.4, 5 mM MgCl2, 150 mM NaCl, 1% Triton X-100, 1% Doxycholate, 2.5 mM DTT; 200 U/ml RNasin, 100 µg /ml CHX, cOmplete, EDTA-free protease inhibitor cocktail (Roche)), and incubated on ice for 10 min with occasional mixing. Lysates were centrifuged at 10,000g for 5 min at 4 °C and the supernatant was carefully removed. For ribosome fractionation: Protein concentrations for lysates were measured by Bradford assay and equal amounts of protein were loaded on a 10– 50% sucrose gradient containing 100 µg/ml CHX, 0.2 mg/ml heparin and 1 mM DTT. Gradients were centrifuged at 41,000 RPM for 2 h at 4 °C in a SW41 rotor (Beckman Coulter Life Sciences) and subsequently fractionated using a Piston Gradient Station (BioComp), measuring absorbance at 254 nm. For standard ribosome profiling: lysis buffer was supplemented with 25U/ml DNaseI- XT (NEB). Ribosome protected fragments were generated by MNase digestion. MNase digestion buffer (300mM NaCl (5M), 50mM Tris PH 7.5, 5mM MgCl2, 0.5% Triton 1mM DTT and 15 mM CaCl2) was added to lysate in a ratio of 1:2 (to a final concentration of CaCl2 of 5mM). Cells were incubated for 20 minutes in 25°C with 1-2 μl of MNase enzyme. MNase was inactivated by adding EGTA to a final concentration of 6.25mM. Lysates were loaded onto 900ul of 1M sucrose cushions (supplemented with CHX and DTT) and centrifuged for 1 hour at 111,000 RPM using a TLA120.2 rotor (Optima max-xp, Beckman). After centrifugation, the pellet was suspended in lysis buffer and RNA was extracted using RNA miniprep kit (Zymo research). RNA fragments were run on a 8M urea 15% acrylamide gel, alongside two lanes containing two oligoribonucleotides each - 26 and 34 nucleotides long, serving as a size indicator. Gel was pre-run for 15 minutes, and electrophoresis lasted 60 minutes in 200 V.

### Click-IT L-homopropargylglycine metabolic labeling and detection

Cells were diluted to ∼30% confluence. The next day cells were washed with PBS and incubated in methionine-free RPMI medium for 30 min. Then L-homopropargylglycine (HPG; Thermo Fisher) was added to the medium according to the manufacturer’s instructions and incubated for 7 minutes. Cells were collected, processed, and labeled with Click-iT™ HPG Alexa Fluor™ 594 Protein Synthesis Assay Kit (cat. C10429) according to the manufacturer’s instructions.

### Human SNORD CRISPR screen

A CRISPR-Cas9 snoRNA-library with 811 sgRNAs targeting known box C/D snoRNA (snoRD) genes, snoRD host genes, as well as positive and nontargeting controls was designed based on the snoRNA-CRISPR library from Pauli et al^8^. The sgRNAs were cloned into the lentiCRISPRv2GFP vector, which was a gift from David Feldser^4^ (Addgene plasmid #82416). For lentiviral propagation, in brief, 293T cells were transfected with the lentiCRISPRv2GFP snoRD library, pVSVG, and psPAX2 using PEI-MAX transfection reagent. At 48 hours after transfection, supernatant containing lentivirus particles was harvested. K562 cells were infected with low multiplicity of infection (=0.3), GFP^+^ cells were sorted 48hrs after transduction, transduction was considered sufficient only if more than 500 cells/sgRNA were sorted. Cells were maintained in culture for 1 week prior to screen initiation. At day 0 at least 1 million cells were collected for genomic DNA extraction or treated with 2nM PMA for 72hrs. Then the cells were stained for CD41/61 (clone A2A9/6) levels and highly differentiated cells were sorted. The sgRNA encoding sequences were amplified by nested polymerase chain reaction, the products were purified from agarose gel. MAGeCK software was used to analyze the results.

### Human SNORD CRISPR screen bioinformatics analysis

First, we trimmed the 5’ adapter sequence TTGTGGAAAGGACGAAACACCG using Cutadapt 1 . The results were then analyzed using MAGeCK 2 software. After sgRNA counting, we retained only the biological replicates where the total number of reads was at least 300 times the number of gRNAs, with a mapped read percentage of at least 60%. Additionally, the number of missing gRNAs was limited to no more than 1%, and the Gini index was kept small (<0.1) to indicate evenness in the count distribution. In the MAGeCK testing step, we combined the four replicates that passed quality control to identify sgRNAs that were positively or negatively selected. The results were visualized using EnhancedVolcano 3, with significance thresholds set at an adjusted p-value of less than 0.05 and absolute fold-change values greater than 0.58.

### Data analysis Ultra-low ribosome profiling and ultra-low RNA sequencing

***Data analysis – Ribo-seq***: ***reference genomes and annotations***. The reference genome and transcript annotations were obtained from Gencode mouse release 24 (GRCm38.p6). The genome was prepared for alignment by masking all tRNA genes and pseudogenes in the chromosome sequences, and including unique mature tRNA genes as artificial chromosomes. These tRNA genes and psueodgenes were identified using tRNAscan-SE (version 2.0.7) using the eukaryotic and vertebrate mitochondrial models. For metagene analyses, a set of canonical transcripts was defined based on the APPRIS annotations, with the longer isoform being selected in cases where multiple primary APPRIS isoforms existed.

***Data analysis – Ribo-seq: read processing, alignment, and quantification*.** Ribo-seq raw reads were processed, aligned, and quantified as described in scRibo-seq^1^. Briefly, the unique molecular identifier (UMI) was extracted from the first 10 bases of read 1 and prefixed to the cell barcode in read 2. Adapter sequences were trimmed from read 1 using cutadapt (version 3.2) with -m 15: -a TGGAATTCTCGGGT. Trimmed reads were aligned to the reference genome using STARsolo (version 2.7.6a) with the following parameters: --sjdbOverhang 50 -- seedSearchStartLmax 10 --alignIntronMax 1000000 --outFilterType BySJout -- alignSJoverhangMin 8 --outFilterScoreMin 0 --outFilterMultimapNmax 1 –chimScoreSeparation 10 --chimScoreMin 20 --chimSegmentMin 15 --outFilterMismatchNmax 5. Aligned reads were deduplicated with UMI-tools (version 1.1.1) using --spliced-is-unique --per-cell --read-length -- no-sort-output, and sorted using sambamba (version 0.8.0). Aligned and deduplicated reads were filtered to select those that aligned to the correct strand, had a trimmed length between 30 and 45 nt, and had a trimmed length equal to the mapped length. Remaining reads were parsed to extract mapping coordinates used for metagene analyses. Count tables report the number of unique filtered reads aligning within any annotated coding sequence for each gene; coding sequences were expanded to include 25 nt upstream and downstream of the start and stop codons, respectively.

***Data analysis – RNA-seq***: Ultra-low raw RNA-seq data were trimmed with Cutadapt (v.4.7). Clean data were aligned to the mouse reference genome GRCm38 by STAR (v.2.7.11b) with default settings. Genes counts were quantified by HTSeq (v.2.0.3). DEGs were determined by DESeq2, FDR<0.05.

For both Ribo-seq and RNA-seq datasets transcript per million (TPM) counts were filtered to remove genes with below 25 in less than 2 samples of at least one cell type. Then remaining zeros were replaced with a random small value off the data’s distribution. Translation efficiency was calculated on these values by log2 transformation and TE=RPF-RNA. Top 50 signature genes were identified by comparing each gene’s expression in each cell type to all other cell types (using Desq2) and choosing the significant (p < 0.05) top 50 most uniquely highly expressed genes of each cell type. sPLS-DA clustering model was performed using R package *‘mixOmics’*, keeping 150 genes.

### Data analysis K562 ribosome profiling

For both ribo-seq and RNA-seq datasets of K562 cells, initial quality control was conducted using FastQC (0.11.9). Then, adapter trimming and length filtering were performed with Cutadapt (3.5). Reads were aligned to the human genome version hg38 using STAR (2.7.10a). For the ribo-seq data, only aligned reads between 16-36 nucleotides were retained using SAMtools (1.16.1). Gene expression counts were quantified using HTSeq (0.13.5) with the ‘union’ mode, specifying ‘CDS’ for ribo-seq and ‘exon’ for RNA-seq. RNA-seq data were normalized to fragments per kilobase of transcript per million mapped reads (FPKM) using transcript length, while ribo-seq data were normalized to coding sequence (CDS) length. Genes with log transcript per million (TPM) counts below 3 in all conditions were filtered out, and differential expression analysis was performed using DESeq2 with false discovery rate (FDR) correction. Next, the datasets were integrated to measure translation efficiency (TE), defined as the ratio of ribo-seq to RNA-seq normalized counts. Significant differential TE was identified as a fold change in logTE between conditions greater than 1.5. Differential codon usage was calculated by R package ‘seqinr’ uco function, each genes codon frequency was calculated and averaged across groups then divided for fold change.

### Data analysis RiboMethSeq

First, we performed quality trimming and adapter clipping using Trimmomatic (v0.39). In RiboMethSeq analysis, only short reads with properly clipped adapters at the 3’-end were retained. Next, we mapped reads to mouse reference rRNA sequences (18S, 28S, 5S, and 5.8S) using Bowtie2, keeping only uniquely mapped reads. Both 5’- and 3’-ends of the fragment were utilized. Methylation levels were quantified by calculating the MethScore for each rRNA position.

### Similarity index

In order to compare the dissimilarity of methylation patterns of known sites between different cell types, we used the Euclidean distance between groups.

